# Widespread non-apoptotic activation of *Drosophila* Caspase-2/9 limits JNK signaling, macrophage proliferation, and growth of wound-like tumors

**DOI:** 10.1101/2020.07.27.223404

**Authors:** Derek Cui Xu, Kenneth M. Yamada, Luis Alberto Baena-Lopez

## Abstract

Resistance to apoptosis due to caspase deregulation is considered one of the main hallmarks of cancer. However, the discovery of novel non-apoptotic caspase functions has revealed unknown intricacies about the interplay between these enzymes and tumor progression. To investigate this biological problem, we capitalized on a *Drosophila* tumor model highly relevant for humans that relies on the concomitant upregulation of EGFR and the JAK/STAT signaling pathway. Our results indicate that widespread non-apoptotic activation of initiator caspases limits JNK signaling and facilitates cell fate commitment in these tumors, thus preventing the overgrowth and exacerbation of malignant features. Intriguingly, these caspase functions are strongly linked to the ability of these enzymes to control the recruitment and subsequent proliferation *in situ* of macrophage-like cells on the tumor. These findings assign novel tumor-suppressor activities to caspases independent of apoptosis, while providing highly relevant molecular details to understanding their diverse contribution during tumor progression.

## Introduction

The ability of caspases to stimulate the apoptosis program is at the forefront of the molecular mechanisms against cancer (Olsson and Zhivotovsky, 2011), and accordingly, defective apoptosis is considered one of the most distinctive features of cancerous cells (Hanahan and Weinberg, 2000, 2011). However, the recent incorporation of non-apoptotic roles to caspases (Aram et al., 2017; Baena-Lopez et al., 2018; Miura, 2012) have added layers of complexity to the interplay between these enzymes and tumor progression (Jager and Zwacka, 2010; Xu et al., 2018). Establishing the biological significance to these novel caspase roles is critical to understand their contribution to tumorigenesis, as well as to develop efficient therapeutic strategies against cancer based on caspase-modulating molecules. Our work in this study has uncovered novel tumor-suppressing caspase functions that rely on their ability to modulate intracellular signaling and the configuration of the tumor microenvironment without causing apoptosis.

Evolutionary conservation of gene function and powerful genetic tools have made *Drosophila melanogaster* an extremely useful model organism to investigate the molecular principles of cancer as well as the strong links with caspases (La Marca and Richardson, 2020; Mirzoyan et al., 2019; Parvy et al., 2018; Perez et al., 2017; Richardson and Portela, 2018; Xu et al., 2018). One commonly studied *Drosophila* tumor model involves a form of the oncogene *Ras*^*V12*^ that constitutively activates the MAP-Kinase (MAPK) transduction pathway independently of upstream tyrosine-kinase receptors such as epidermal growth factor receptor (EGFR) (Richardson and Portela, 2018). These tumor models also often incorporate secondary mutations in essential genes that normally ensure apicobasal polarity of cells, such as *discs-large* or *scribble* (Brumby and Richardson, 2003; Pagliarini and Xu, 2003; Richardson and Portela, 2018). Although these oncogenic mutations alone display limited tumorigenicity, their synergetic combination exacerbates the malignancy of transformed cells in both *Drosophila* and mammalian models (Dow et al., 2008; Kajita and Fujita, 2015; Mirzoyan et al., 2019; Richardson and Portela, 2018; Zhan et al., 2008). Intriguingly, the condition that often underlies this enhanced oncogenic transformation is the persistent hyperactivation of the c-Jun N-terminal Kinase (JNK) pathway (Beira et al., 2018; La Marca and Richardson, 2020; Pinal et al., 2019).

Under physiological conditions, transient JNK activation facilitates resetting of cell identity within damaged *Drosophila* tissues and helps to drive the process of wound healing through cell migration and proliferation (Ahmed-de-Prado et al., 2018; Bergantinos et al., 2010; Pinal et al., 2019; Santabarbara-Ruiz et al., 2015). Furthermore, it can trigger the caspase-pathway to eliminate undesired cells through apoptosis (La Marca and Richardson, 2020; Pinal et al., 2019). Conversely, caspase activation can also enhance JNK signaling through molecular feedback loops that remain poorly characterized (Pinal et al., 2019). Importantly, deregulation of these feedback loops between caspases and JNK signaling can amplify the malignancy of transformed cells through mechanisms that are starting to be elucidated (Ahmed-de-Prado et al., 2018; Berthenet et al., 2020; Perez et al., 2017; Pinal et al., 2019; Pinal et al., 2018).

We have capitalized on a *Drosophila* model of cooperative oncogenic transformation that relies on simultaneous overactivation of the **E**GFR and **J**AK/**S**TAT signaling pathways (we hereafter refer to this model as **EJS**) (Herranz et al., 2012) to investigate the functional diversity of caspases during tumor progression. EJS tumors are a particularly attractive model from an apoptotic perspective, since their signaling profile can provide potent pro-survival cues that buffer the activation of the apoptotic program. Specifically, whereas EGFR signaling can negatively regulate the activity of pro-apoptotic factors such as Hid (Bergmann et al., 1998), JAK/STAT activation can induce the expression of apoptosis inhibitors such as *Diap-1* in *Drosophila* (Betz et al., 2008; Recasens-Alvarez et al., 2017) or *Bcl-2* proteins in mammals (Fujio et al., 1997). From a cancer perspective, the oncogenic cooperation between EGFR and JAK-STAT is also highly relevant since it has been shown to transform primary BJ fibroblasts in cell culture (Herranz et al., 2012) and sustain tumor growth and malignant transformation of various human cancers (Andl et al., 2004; Quesnelle et al., 2007; Sun et al., 2016).

Combining the EJS tumor model and a highly sensitive caspase sensor developed in our laboratory (Baena-Lopez et al., 2018), we provide seminal evidence indicating that virtually all of the cells forming a tumor can have non-apoptotic caspase activation. Furthermore, we show that such non-lethal caspase activity is critical for limiting JNK signaling and the exacerbation of malignant features in EJS tumors such as excess growth and poor cell differentiation. Strikingly, we have correlated these novel caspase effects with the non-autonomous regulation of features of *Drosophila* tumor-associated macrophages (DTAMs). These findings confer non-apoptotic tumor suppressor properties to caspases, while offering new molecular leads to understanding the complex interplay between JNK signaling, caspases, and DTAMs during tumor progression.

## Results

### EJS tumors have widespread non-apoptotic caspase activity

As previously shown, EJS tumors can be induced in *Drosophila* wing discs with spatial and temporal precision using the *ap-Gal4* driver and a temperature sensitive Gal80 protein (Gal80^ts^) (Herranz et al., 2012). The EJS genetic configuration allows the efficient oncogenic transformation of the dorsal cells forming the wing discs by shifting larvae from 18°C to 29°C (Figure 1A). To investigate the dynamics of caspase activation during tumor progression, we combined the EJS tumor model with a sensitive Drice-based caspase sensor (DBS-S-QF), which specifically reports on *Drosophila* initiator caspase activation (Baena-Lopez et al., 2018). This tool can transiently label caspase-activating cells with a short-lived fluorescent protein (Tomato-HA) as well as with a permanent cellular marker (beta-galactosidase, β-gal), thus providing a history of caspase activation (Baena-Lopez et al., 2018). The presence of permanently labelled cells (β-gal) without signs of ongoing caspase activation (Tomato-HA) or cell death (e.g. nuclear fragmentation) represents an unambiguous demonstration of non-apoptotic caspase activity, as these cells either themselves activated caspases in the past but did not complete apoptosis, or are directly descended from such cells (Baena-Lopez et al., 2018). While a modest fraction of cells were labeled with either the transient (HA immunostaining) or permanent caspase markers (β-gal) in wild-type wing discs (Baena-Lopez et al., 2018) (Figure 1B,C), a large proportion of cells were decorated by anti-HA in EJS tumors, and virtually 100% of the transformed cells expressed β-gal (Figure 1D); only a few residual wild-type cells without *ap-Gal4* and EJS expression remained negative for β-gal (Figure 1E). These results demonstrate for the first time that caspases can be widely activated within tumors without causing the elimination of transformed cells through apoptosis. Moreover, caspase-activating cells can remain proliferative as part of the tumor mass (Figure 1D).

**Figure 1.**
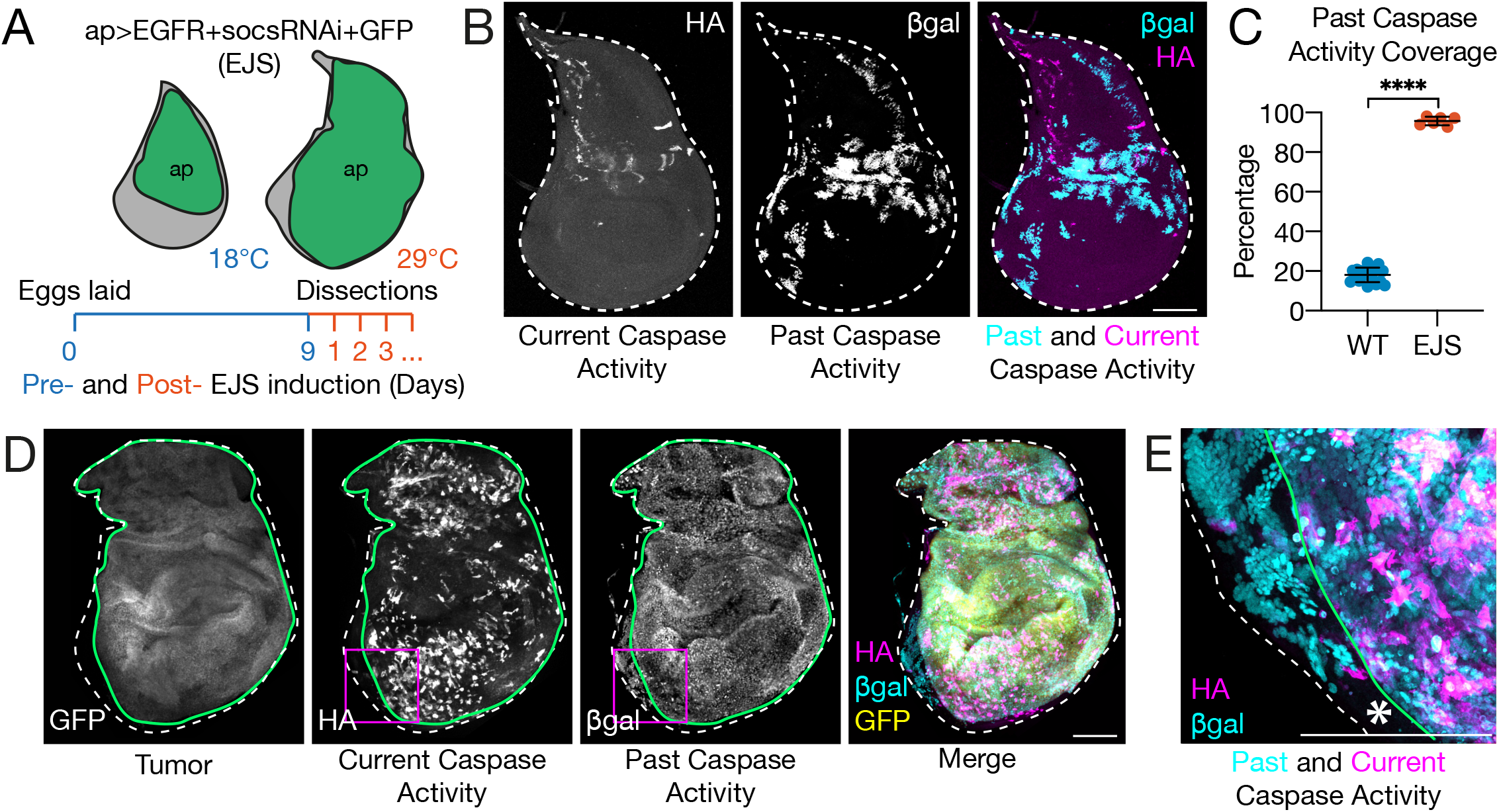
Non-apoptotic caspase activation in EJS tumors. **(A)** Schematic showing the experimental design for inducing tumors based on EGFR and JAK/STAT **(EJS)**upregulation in imaginal wing discs. Dorsal cells of the wing discs expressing *apterous-Gal4* (ap, green) express *UAS-GFP*, *UAS-EGFR*, and *UAS-socs36E-RNAi* transgenes after larvae are temperature shifted from 18°C to 29°C. Ubiquitously expressed *Tubulin-Gal80*^*ts*^ prevents transgene expression and thus tumor induction in larvae for 9 days after egg laying while at 18°C. After larvae were transferred to 29°C to induce tumor formation, dissections occurred at specific time points 1 – 5 days after temperature shift. **(B)** Lineage tracing of caspase-activating cells in wild-type wing discs using the DBS-S-QF sensor. Representative maximum projected confocal images showing cells in third instar wing discs labeled for past caspase activity (gray and cyan) and current caspase activity (gray and magenta) in the same fashion as in **(B)**. The entire wing disc is outline (white dashed outline) using a DAPI stain as a reference (not shown). Scale bar: 150μm. **(C)** Quantification of the area of wing discs composed of cells expressing the marker for past caspase activity as a percentage of total wing disc area in wild-type wing discs and EJS tumors. Mean ±SD are plotted. Statistical significance was determined using an unpaired Student’s t-test; **** p<0.0001. Numbers of wing discs analyzed for wild-type discs: 18; for EJS tumors: 6. **(D)** Lineage tracing of caspase-activating cells in EJS tumors using the DBS-S-QF sensor. Representative maximum projected confocal images showing EJS tumors after 3 days of EJS induction labeled with GFP expression (gray and yellow). Cells with past caspase activity are permanently labeled with LacZ expression immunostained with anti-beta-galactosidase (gray and cyan). Cells with current caspase activity express a *QUAS-mTdTomato-HA* transgene (gray and magenta). The area of the wing disc composed of EJS tumor cells is outlined (green outline) using GFP expression as a reference, while the entire wing disc is outlined (white dashed outline) using a DAPI stain as reference (not shown). Scale bar: 100μm. **(E)** Representative maximum projected confocal image at higher resolution indicating the overlap in EJS tumors of cells with caspase activity at the time of dissection (anti-HA, magenta) and in the past (anti-beta-galactosidase, cyan) within the region of the wing disc indicated by the magenta squares. Notice the presence of cells expressing only the cellular marker corresponding to past caspase activity (cyan) and the small fraction of cells not showing any caspase activity (asterisk). Scale bar: 100μm. Full genotype description can be found in MM.

### *Drosophila* Caspase-2/9 activation compromises tumor growth independently of apoptosis

Because the DBS-S-QF caspase sensor was designed to detect the activity of upstream initiator caspases (Baena-Lopez et al., 2018), and the major initiator caspase in *Drosophila* outside of the immune system is *Dronc* (*Drosophila* ortholog of mammalian Caspase-2/9), we investigated the contribution of this caspase member to the progression of EJS tumors. Intriguingly, the depletion of *Dronc* expression using a well-characterized RNAi transgene (Leulier et al., 2006) did not noticeably alter tumor growth before two days had passed following tumor induction (Figure 2A); however, by the third day, *Dronc* deficiency triggered a prominent tumor expansion that only increased over time (Figures 2A-C). Confirming the specificity of these phenotypes, equivalent results were obtained with a conditional *Dronc*^*KO*^ allele (Arthurton et al., 2019) (Figure 2D). These results indicated a tumor-suppressing role for *Dronc* in EJS tumors.

**Figure 2.**
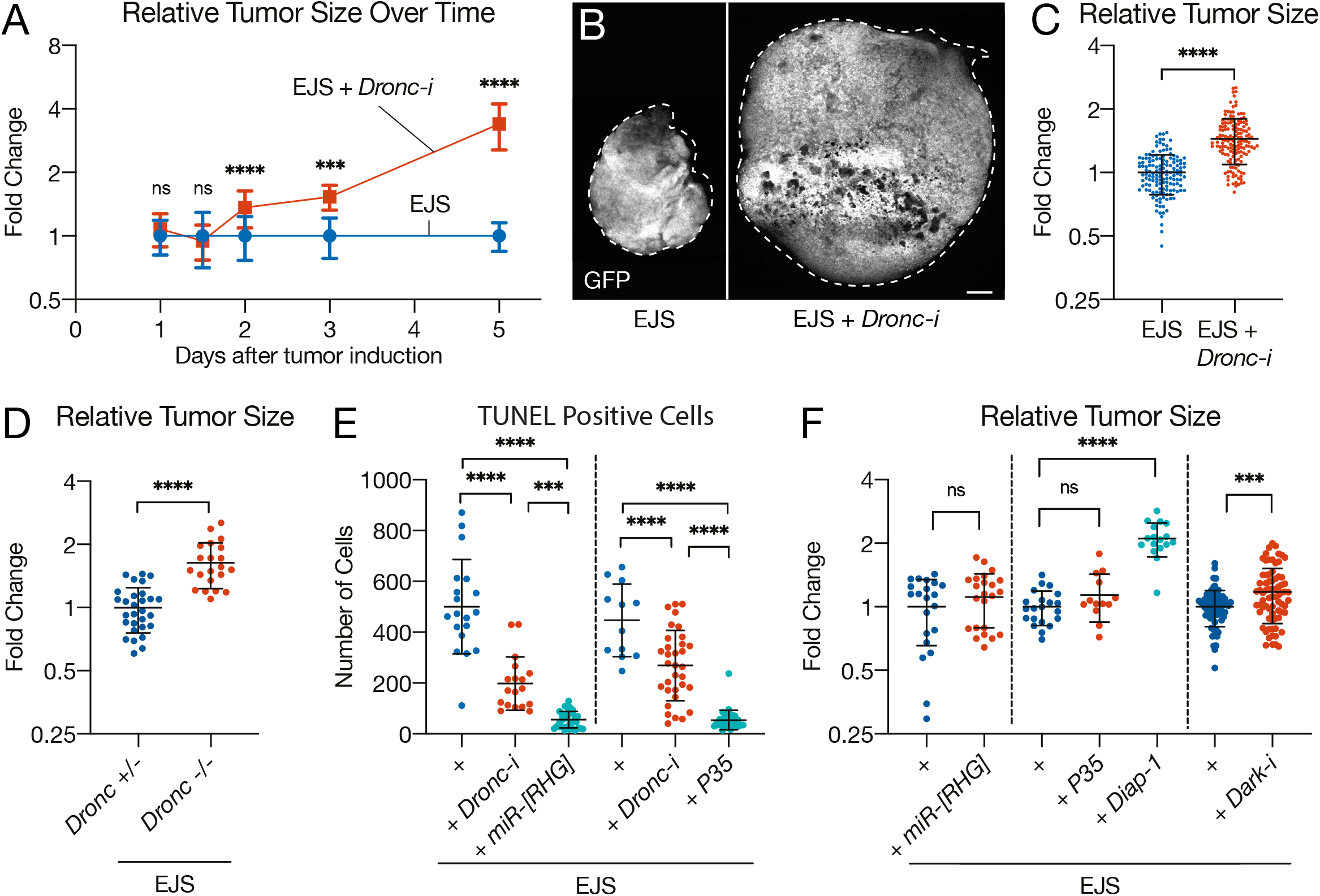
Non-apoptotic activity of initiator caspases restricts tumor growth and size. **(A)** Graph indicating the relative size of EJS tumors (blue) and EJS tumors expressing a Dronc-RNAi construct (*Dronc*-i) (orange) over time after tumor induction. Plotted are the mean ±SD of relative tumor sizes at each time point. Notice the differences in size two days after tumor induction. Control tumors were used for normalization. One-way ANOVA with Tukey’s multiple comparisons tests was used to determine statistical significance at each time point; ns – not significant p>0.05, *** p=0.001, **** p<0.0001. Numbers of wing discs analyzed for each time point are as follows (**Day**: n [EJS]; n [EJS + *Dronc*-i]): (**1**: 39; 36), (**1.5**: 21; 38), (**2**: 29; 35), (**3**: 18; 7), (**5**: 12; 12). **(B)** Representative confocal images of control EJS and EJS + *UAS-Dronc-RNAi* tumors after 5 days of EJS induction (GFP, gray). The entire wing disc is outlined (white dashed outline) using a DAPI stain (not shown). Full genotype description can be found in MM. Scale bar: 100μm. **(C)** Relative sizes of control EJS and EJS + *UAS-Dronc-RNAi* (EJS + *Dronc-i*) tumors after 3 days of EJS induction. Control tumors were used for normalization. Statistical significance was determined by using an unpaired Student’s t-test; **** p<0.0001. Numbers of wing discs analyzed for EJS tumors: 145; for EJS + *Dronc*-i tumors: 154. **(D)** Relative sizes of control (*Dronc*^+/−^) and EJS *Dronc*^−/−^ (*Dronc*^−/−^) tumors either heterozygous or homozygous mutant for *Dronc*, respectively (full genotype in MM). Control tumors were used for normalization. Statistical significance was determined by an unpaired Student’s t-test; ****p<0.0001. Numbers of wing discs analyzed for control tumors: 29; for EJS *Dronc*^−/−^ tumors: 21. **(E)** Quantification of apoptosis detected by TUNEL-positive cells in control EJS (+), EJS + *UAS-Dronc-RNAi* (+ *Dronc-RNAi*), EJS + *UAS-miR[RHG]* (+ *miR[RHG]*), and EJS + *UAS-P35* (+ *P35*) tumors (full genotype in MM). Statistical significance was determined by one-way ANOVA with Tukey’s multiple comparisons test; *** p=0.0001, **** p<0.0001. Numbers of wing discs analyzed for EJS tumors: 19 and 12; for EJS + *Dronc*-i tumors: 18 and 33; for EJS + miR[RHG] tumors: 33; for EJS + P35 tumors: 34. **(F)** Relative sizes of control EJS (+), EJS + *UAS-miR[RHG]* (+ *miR[RHG]*), EJS + *UAS-P35* (+ *P35*), EJS + *UAS-Diap-1* (+ *Diap-1*), and EJS + *UAS-Dark-RNAi* (+ *Dark-i*) tumors (full genotype in MM). Statistical significance was determined by an unpaired Student’s t-test for (+ *miR[RHG]*) and (+ *Dark-i*), and one-way ANOVA with Tukey’s multiple comparisons tests for (+ *P35*) and (+ *Diap-1*). ns – not significant p>0.05, *** p=0.0004, **** p<0.0001. For all graphs, mean ±SD are also plotted. Numbers of wing discs analyzed for EJS tumors: 21, 21, and 65; for EJS + miR[RHG] tumors: 23; for EJS + P35 tumors: 13; for EJS + Diap-1: 17.

Since the tumor-suppressing ability of caspases has conventionally been linked to the pro-apoptotic role of these enzymes (Hanahan and Weinberg, 2000, 2011), we investigated whether the increase in tumor size could be explained by the accumulation of apoptosis-resistant cells. To this end, we inhibited the apoptotic pathway by either silencing the expression of upstream pro-apoptotic factors Reaper, Hid, and Grim (RHG) (Siegrist et al., 2010), or by blocking the activity of downstream effector caspases through overexpression of P35 (Hay et al., 1994). As indicated by TUNEL assays, these genetic manipulations effectively impeded apoptosis (Figure 2E); however, they failed to replicate the overgrowth phenotypes caused by *Dronc*-deficiency (Figure 2F). On the contrary, the inhibition of *Dronc* through overexpression of *Drosophila* inhibitor of apoptosis Diap-1 mimicked the *Dronc*-deficient overgrowth phenotypes (Figure 2F). To define the origin of Dronc activation in EJS tumors, we compromised the expression of *Dark* (orthologous to *Apaf-1* in mammals). Upon binding, Dark and Dronc form a protein complex termed the apoptosome that facilitates the efficient activation of Dronc during apoptosis (Hay et al., 2004). The overexpression of a *Dark-RNAi* construct in EJS tumors caused a less-penetrant but consistent tumor overgrowth (Figure 2F), thus suggesting the involvement of this apoptotic component in the activation of Dronc. These results in combination with those obtained with the DBS-S-QF sensor strongly indicated the non-apoptotic nature of Dronc activation in EJS tumors, and uncoupled the *Dronc* loss-of-function phenotypes from defective apoptosis.

### Non-apoptotic Dronc activity limits proliferation and cell size in EJS tumors

To determine which factors could explain the increased tumor size in *Dronc*-deficient tumors, we assessed the amount of cell proliferation in this genetic condition using the cell cycle markers phospho-histone-H3 (PH3) and 5-ethynyl-2′-deoxyuridine (EdU). Immunostainings showed an upregulation of the cell proliferation markers in caspase-deficient EJS tumors (Figure 3A-C). These results suggested that *Dronc* limits cell proliferation within EJS tumors. In addition, using nuclear size as a proxy for cell size (Cantwell and Nurse, 2019), we observed significant cell enlargement and decreased cell density within *Dronc*-deficient tumor cells (Figure 3D,E, Figure S1). These experiments suggested that caspase deficiency promotes tumor overgrowth by increasing both the number and size of transformed cells.

**Figure 3.**
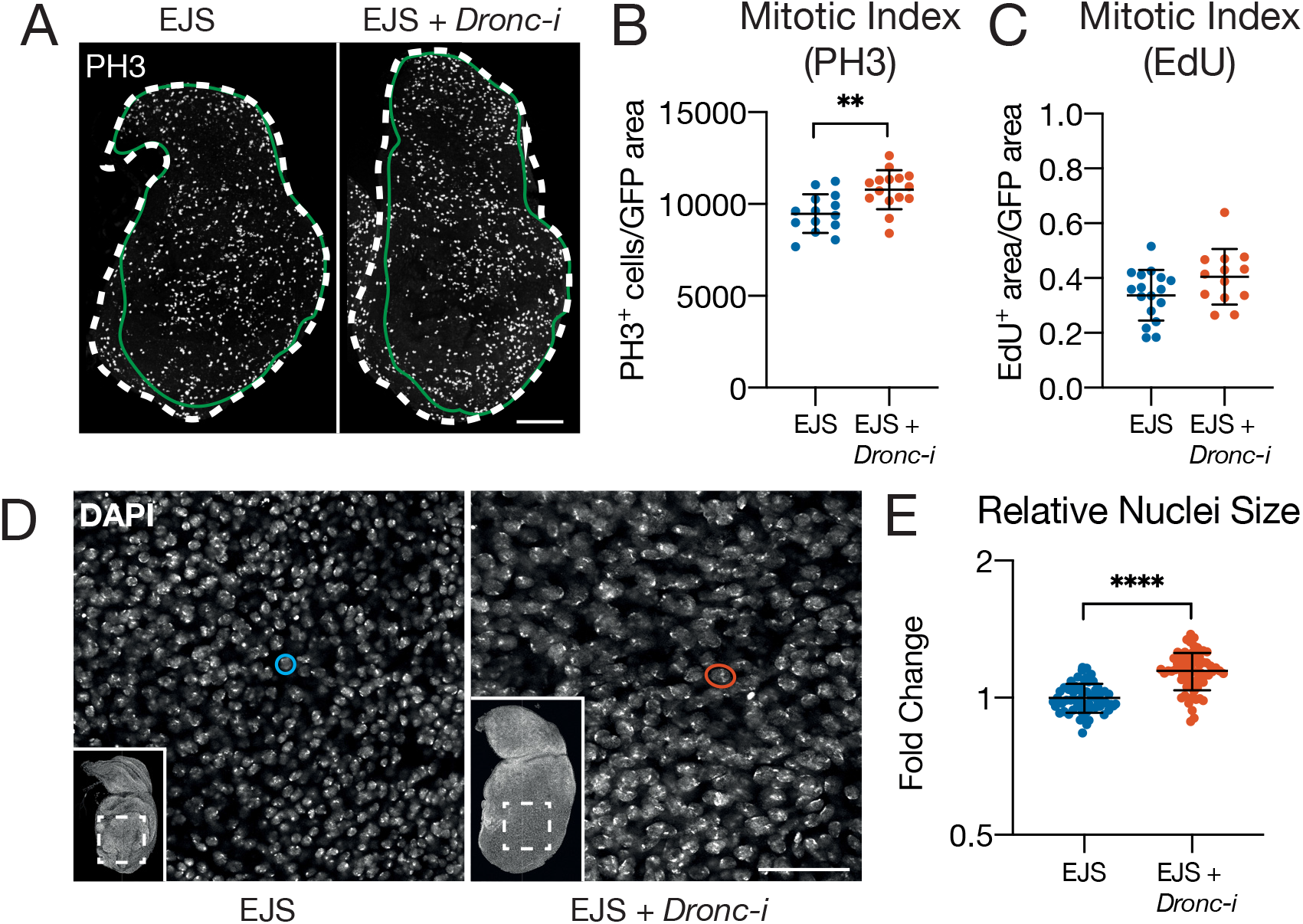
Non-apoptotic activity of initiator caspases restricts proliferation and cell size in EJS tumors. **(A)** Representative confocal images of control (EJS) and EJS + *UAS-Dronc-RNAi* (EJS + *Dronc*-i) tumors after 1 day of EJS induction showing phospho-Histone H3 (PH3) immunostaining (gray). Outline of wing disc (dashed white) and tumor (green) obtained by tracing DAPI and GFP, respectively. Scale bar: 100μm. **(B,C)** Quantification of PH3^+^ (B) and EdU^+^ (C) cells in control (EJS) and EJS + UAS-Dronc-RNAi (EJS + *Dronc*-i) tumors. Statistical significance was determined by an unpaired Student’s t-test; p>0.05, ** p=0.0025. Numbers of wing discs analyzed for control tumors: 14 in (B) and 17 in (C); for EJS + *Dronc*-i tumors: 15 in (B) and 13 in (C). Mean ±SD are also plotted. **(D)** Higher magnification (60x) confocal image of nuclei stained with DAPI in control (EJS) and EJS + *UAS-Dronc-RNAi* (EJS + *Dronc*-i) tumors after 3 days of EJS induction. Inset depicts the entire tumorous wing disc with the dashed outline indicating the main magnification image. Example nuclei for size comparison are circled in blue (EJS) and orange (EJS + *Dronc*-i). Scale bar: 50μm. **(E)** Relative sizes of nuclei in control (EJS) and EJS + *UAS-Dronc-RNAi* (EJS + *Dronc-i*) tumors after 3 days of EJS induction. Nuclei in control EJS tumors were used for normalization. Mean±SD are also plotted. Statistical analysis performed by Student’s t-test. ****P<0.0001. Numbers of wing discs analyzed for control EJS tumors: 59; for EJS + *Dronc*-i: 65.

### Caspase-dependent inhibition of JNK signaling limits neoplastic transformation

Because JNK activation often sits at the origin of malignant transformation (Beira et al., 2018; La Marca and Richardson, 2020; Wu et al., 2019) and promotes caspase activation (Dhanasekaran and Reddy, 2017; Pinal et al., 2019; Pinal et al., 2018), we explored the potential interplay between this signaling pathway and caspases in EJS tumors.

To determine the JNK pathway activation in EJS tumors, we evaluated the expression of the universal JNK target gene MMP1 (Uhlirova and Bohmann, 2006) and a synthetic transcriptional reporter (Tre-RFP) (Chatterjee and Bohmann, 2012). EJS tumors showed a consistent upregulation of both JNK readouts, which in turn was significantly increased in *Dronc*-deficient conditions (Figures 4A-D). Confirming these effects, qPCR experiments showed robust transcriptional upregulation of Unpaired (Upd) ligands in *Dronc*-deficient transformed cells (Figure 4E). Upd ligands are additional JNK target genes induced in tumors as well as during regeneration (Ahmed-de-Prado et al., 2018; Jiang et al., 2009; La Marca and Richardson, 2020; Pastor-Pareja et al., 2008; Santabarbara-Ruiz et al., 2015; Worley et al., 2018; Wu et al., 2010). Collectively, these findings strongly suggest that Dronc limits JNK signaling in EJS tumors.

**Figure 4.**
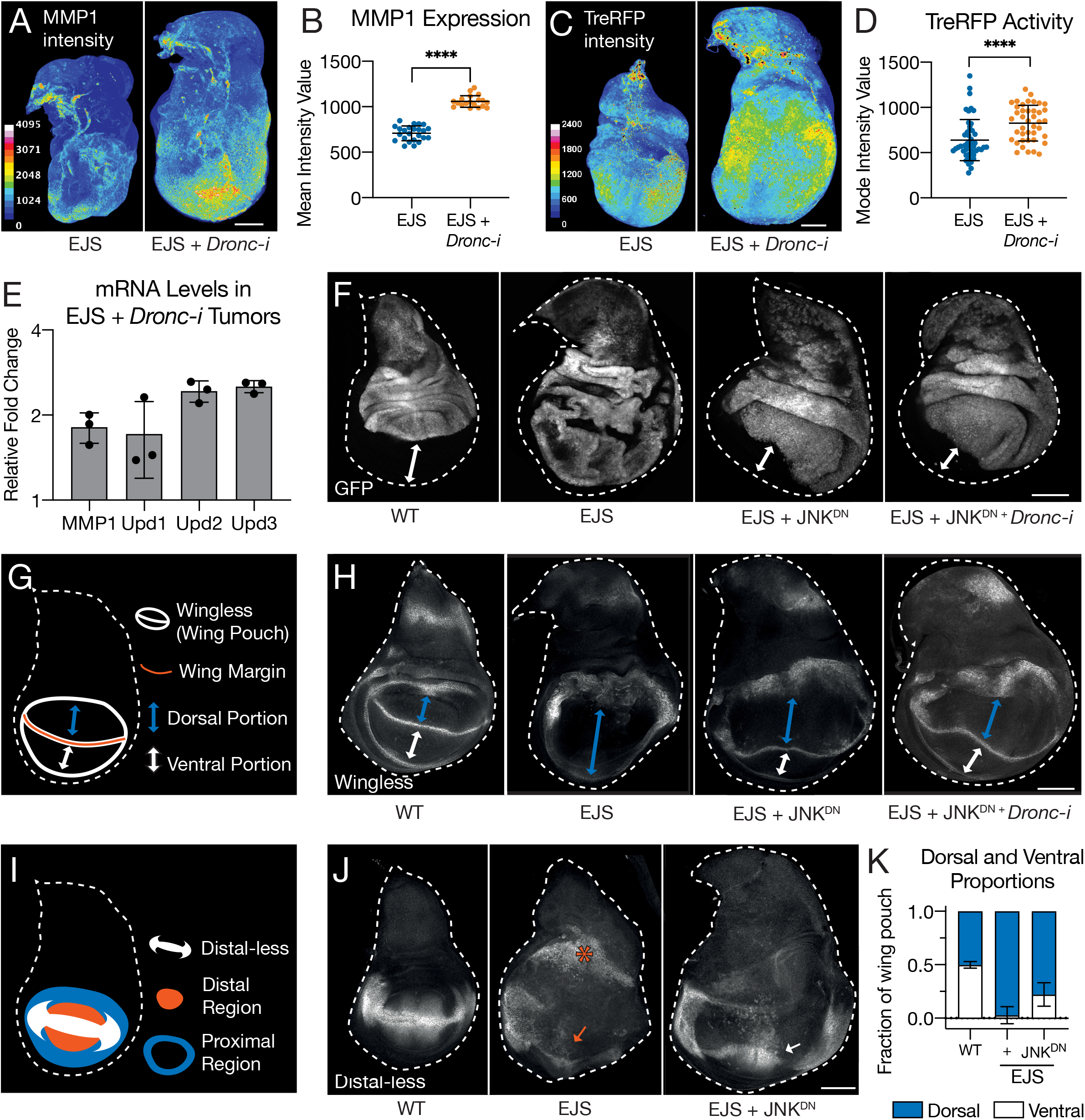
Non-apoptotic activity of Dronc limits JNK signaling responsible for malignant features in open-wound-like EJS tumors. **(A)** Representative confocal images of control (EJS) and EJS + *UAS-Dronc-RNAi* (EJS + *Dronc*-i) tumors after 3 days of EJS induction stained with anti-MMP1, false colored to visualize intensity of MMP1 staining. The intensity scale bar shows the full range of pixel intensities from 0 – 4095. Scale bar: 100μm. **(B)** Quantification of mean MMP1 staining intensity in control (EJS) and EJS + *UAS-Dronc-RNAi* (EJS + *Dronc*-i) tumors after 3 days of EJS induction. Mean ±SD are also plotted. Statistical significance was determined by an unpaired Student’s t-test; ****p<0.0001. Numbers of wing discs analyzed for control EJS tumors: 24; for EJS + *Dronc*-i tumors: 19. **(C)** Representative confocal images of Tre-RFP expression in control (EJS) and EJS + *UAS-Dronc-RNAi* (EJS + *Dronc*-i) tumors after 3 days of EJS induction, false colored to visualize intensity of Tre-RFP expression. The intensity scale bar has been adjusted to show the range of pixel intensities from 0 – 2400 out of the full 4095, to better visualize the differences in intensity. **(D)** Quantification of the mode of Tre-RFP intensity in control (EJS) and EJS + *UAS-Dronc-RNAi* (EJS + *Dronc*-i) tumors after 3 days of EJS induction. Mean ±SD are also plotted. Statistical significance was determined by an unpaired Student’s t-test; ****p<0.0001. Numbers of wing discs analyzed for control EJS tumors: 48; for EJS + *Dronc*-i tumors: 41. **(E)** Relative levels of MMP1, Upd1, Upd2, and Upd3 mRNAs in EJS + *UAS-Dronc-RNAi* (EJS + *Dronc*-i) tumors after 3 days of tumor induction compared to control EJS tumors measured by RT-qPCR. Bars represent the mean of three independent replicates measured, and SDs are also plotted. **(F)** Representative confocal images of wild-type wing discs and control (EJS), EJS+UAS-*bsk*^DN^ (EJS+JNK^DN^), EJS+UAS-*bsk*^DN^ + UAS-*Dronc*-RNAi (EJS+JNK^DN^+*Dronc*-i) tumors after 1.5 days of EJS induction, as EJS + bsk^DN^ tumors seldom progressed past two days post EJS induction due to larval pupariation. GFP (gray) labels the *ap-*expression cells in the dorsal component of the wing discs. Dashed white outline of wing discs obtained by tracing DAPI staining (not shown). White double-headed arrows indicate the ventral portion of the wing discs. Note the absence of a ventral portion in EJS tumors, which is rescued by JNK inhibition (+ JNK^DN^). **(G)** Simplified diagram indicating the expression pattern of Wingless in imaginal wing discs outlining the presumptive wing blade (wing pouch). The wing margin is indicated in orange, which divides the wing pouch into dorsal and ventral portions, indicated by blue and white double-headed arrows, respectively. **(H)** Representative confocal images of Wingless immunostaining (gray) in wild-type wing discs and control (EJS), EJS+UAS-*bsk*^DN^ (EJS+JNK^DN^), EJS+UAS-*bsk*^DN^ + UAS-*Dronc*-RNAi (EJS+JNK^DN^+*Dronc*-i) tumors after 1.5 days of EJS induction. Dashed white outline of wing discs obtained by tracing DAPI staining (not shown). In the same manner as in **(G)**, blue and white double-headed arrows refer to the dorsal and ventral portions of the presumptive wing. Notice the lack of a ventral portion in EJS tumors. **(I)** Simplified diagram indicating the expression pattern of Distal-less in imaginal wing discs. The distal regions of the presumptive wing are marked with orange, while the proximal regions are marked with blue. **(J)** Representative confocal images of Distal-less (Dll) immunostaining (gray) in wild-type wing discs and control (EJS), EJS+UAS-*bsk*^DN^ (EJS+JNK^DN^), EJS+UAS-*bsk*^DN^ + UAS-*Dronc*-RNAi (EJS+JNK^DN^+*Dronc*-i) tumors after 1.5 days of EJS induction. Dashed white outline of wing discs obtained by tracing DAPI staining (not shown). For EJS, the orange arrow indicates downregulated Dll in the distal region of the wing pouch, while the orange asterisk indicates exogenous Dll expression outside the wing pouch. Notice the return of Dll expression in the distal region in EJS + JNK^DN^ tumors (white arrow). **(K)** Quantifications of the dorsal and ventral portions of the wing pouch as fractions of the wing pouch region in WT, EJS, and EJS + JNK^DN^ tumors after 1.5 days of EJS induction. Wingless was used to identify the wing pouch and margin, as in (G) and (H). Numbers of wing discs analyzed for wild-type discs: 11; control EJS tumors: 44; for EJS + *Dronc*-i tumors: 26.

To determine the role of JNK activation in EJS tumors, we overexpressed a widely used JNK dominant-negative construct *basket*^*DN*^ (*bsk*^*DN*^, referred to as *JNK*^*DN*^). The overexpression of this construct abolished the detection of MMP1 immunoreactivity (Figure S2) and the distinctive morphological disorganization of EJS tumors (Figures 4F). To consolidate the latter finding, we investigated the expression pattern of two cell identity genes, *Distal-less* (*Dll*) and *wingless* (*wg*) (Panganiban, 2000; Swarup and Verheyen, 2012), in the presumptive wing cells (Figure 4G-J). Under wild-type conditions, these two genes show a well-defined expression pattern in the presumptive wing region (Figure 4G,I) that was severely disrupted in EJS tumors (Figure 4H,J). In addition, as previously described (Herranz et al., 2012), EJS transformed cells also compromised the growth and patterning of adjacent wild-type cells not expressing the *apterous-Gal4* driver (ventral cells; Figure 4H,J). Importantly, both the non- and the cell-autonomous phenotypes were largely rescued after blocking JNK activity (Figure 4H,J). Furthermore, this phenotypic suppression remained after blocking JNK signaling and caspase expression at the same time (Figure 4F,H). Together, our observations indicated that caspases limit JNK activation and caspase deficiency requires JNK signaling to exacerbate malignant features of EJS tumors. Independently, they confirmed the ability of JNK signaling to maintain wing cells in an undifferentiated stem-like state, both in tumor models (including EJS tumors) and during regeneration (Ahmed-de-Prado et al., 2018; Beira et al., 2018; Katsuyama and Paro, 2011; Santabarbara-Ruiz et al., 2015; Worley et al., 2018).

### JNK pathway activation is independent of ROS and its conventional ligand receptors in EJS tumors

Because the caspase-deficient EJS phenotypes were strongly correlated with JNK upregulation, we next investigated the potential origin of this JNK activation. Recently, it has been shown that the caspase-dependent release of ROS can attract hemocytes (*Drosophila* macrophages) towards tumor cells, which subsequently stimulate JNK signaling and promote tumor growth through the release of Eiger, a JNK ligand (Fogarty et al., 2016; Perez et al., 2017). Therefore, we assessed the levels of ROS in our experimental conditions using the dihydroethidium (DHE) indicator. While DHE labelling was readily detected in EJS tumors, it disappeared in *Dronc*-deficient EJS tumors (Figure S3A,B). This result confirmed the previously described caspase-dependent generation of ROS in tumors (Perez et al., 2017). To examine the functional contribution of ROS to the EJS features, we compromised the production of either extracellular or intracellular reactive oxygen species, by either silencing the expression of Duox (Ha et al., 2009) or overexpressing Catalase (Ha et al., 2005), respectively. The downregulation of Duox expression suppressed DHE staining in EJS tumors but the overexpression of Catalase did not cause noticeable effects (Figure S3C). These results suggested that the main source of ROS in EJS tumors was intracellular. However, unexpectedly, none of these genetic manipulations prevented the hyperplasia in EJS tumors, nor the recruitment of hemocytes (Figure S3C). These results suggested that the caspase-dependent production of ROS is largely dispensable for sustaining JNK signaling and EJS tumour features. Since hemocytes were still recruited towards EJS tumors through ROS-independent mechanisms, we investigated whether the secretion of JNK ligands (e.g. Eiger/TNF-α) from hemocytes could be responsible for the activation of this pathway in EJS tumors (Perez et al., 2017). To this end, we simultaneously downregulated the expression of the both recognized JNK receptors, *grindelwald* (*grnd*) and *wengen* (*wgn*) in EJS tumors (Andersen et al., 2015; Kanda et al., 2002), thus preventing a potential functional redundancy between both receptors. However, this genetic manipulation failed to compromise the growth of EJS tumors (Figure S3D). These data suggested that the activation of JNK signaling is largely independent of the conventional JNK ligands and receptors as well as the generation of ROS. In addition, ROS generation in EJS tumors is strikingly dispensable for the recruitment of hemocytes.

### Caspase activity and JNK signaling modulate the recruitment and subsequent proliferation *in situ* of tumor-associated hemocytes

Our previous experiments revealed the presence of a large number of hemocytes adhered to EJS tumors that can potentially release soluble factors contributing to tumor growth (Agaisse et al., 2003; Cordero et al., 2010; Jiang et al., 2009; Perez et al., 2017; Woodcock et al., 2015). Therefore, we explored the potential role of caspases in the recruitment of hemocytes to the tumor, i.e. of *Drosophila* tumor-associated macrophages (DTAMs). Intriguingly, *Dronc* downregulation increased the number of DTAMs attached to EJS tumors (Figure 5A,B). Furthermore, this increase in DTAM number appeared soon after tumor initiation (1 day after temperature shift) and before differences in size between caspase-deficient EJS tumors and their corresponding tumor controls were detected (after 2 days EJS induction). This result suggested that caspases can limit the initial recruitment of DTAMs. Intriguingly, we also noticed that the difference in the number of DTAMs increased further two days after tumor initiation (Figure 5B), suggesting the contribution of additional factors beyond cell recruitment to the increased number of DTAMs. To evaluate the role of cell proliferation in this new caspase-dependent phenotype, we looked into the proliferative state of the recruited DTAMs using EdU incorporation. These experiments revealed an unexpected labeling of DTAMs with EdU (Figure 5C), since it is the first time to the best of our knowledge that *in situ* hemocyte proliferation has been reported. While *Dronc*-deficient EJS tumors had increased numbers of DTAMs overall, there was initially no correlation between EdU^+^ DTAMs and the total number of DTAMs (Figure 5D); however, two days after EJS induction, the number of Edu^+^ DTAMs and the total number of DTAMs were strongly correlated in EJS tumors (Figure 5E). Furthermore, this correlation was significantly enhanced compared to control EJS tumors (Figure 5E). These results indicated that caspase activation limits both the initial recruitment and subsequent proliferation of DTAMs on EJS tumors.

**Figure 5.**
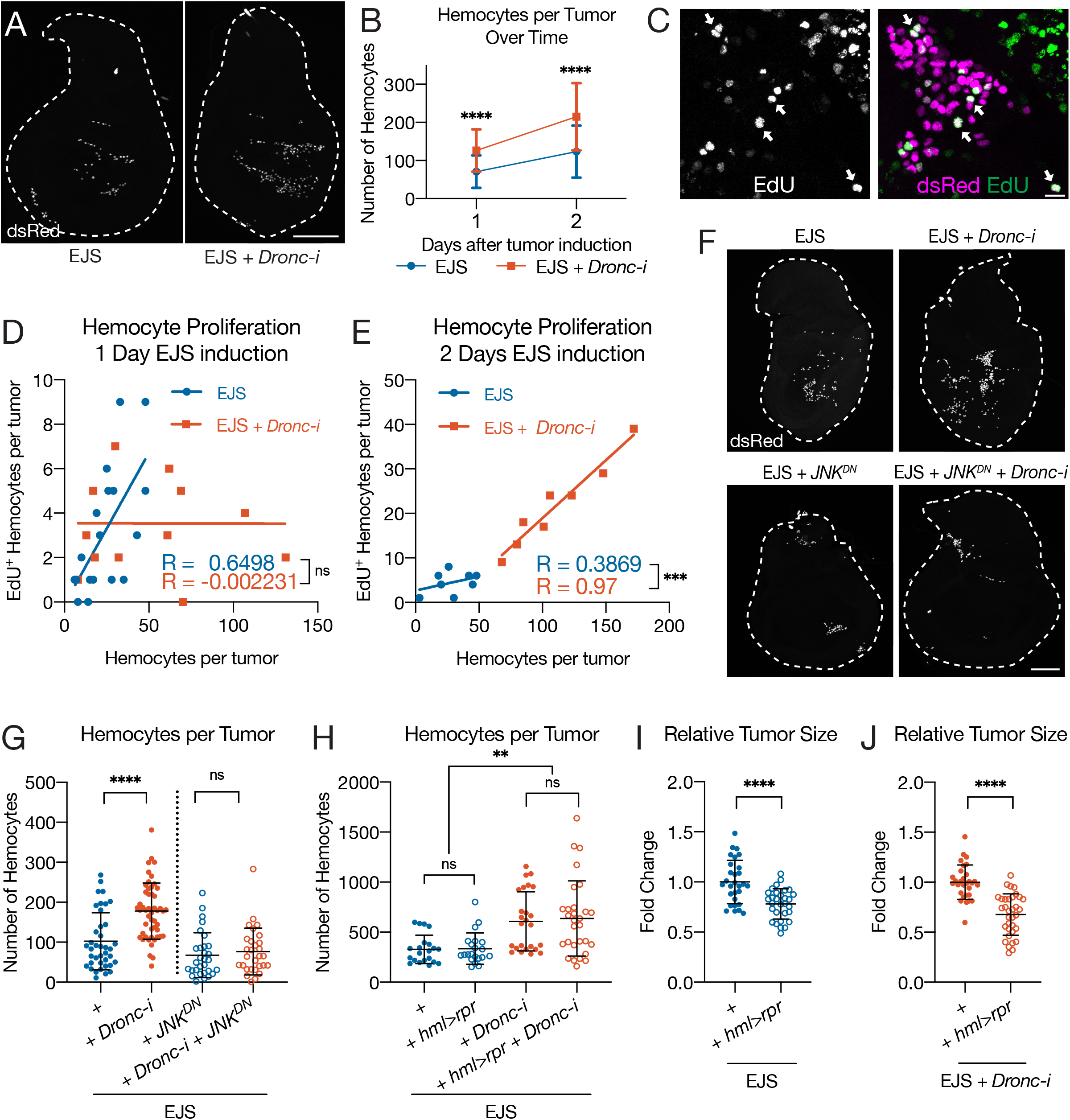
Non-apoptotic caspase activity in EJS tumors limits immune cell recruitment and proliferation. **(A)** Representative confocal images of control (EJS) and EJS + UAS-*Dronc*-RNAi (EJS + *Dronc*-i) tumors after 1 day of EJS induction with hemocytes labeled with *hemolectin-*nls-dsRed *(hml-* dsRed). Dashed white outline of wing discs obtained by tracing DAPI staining (not shown). Scale bar: 100μm. **(B)** Number of hemocytes per tumor in control (EJS, blue) and EJS + UAS-*Dronc*-RNAi (EJS + *Dronc*-i, orange) after 1 and 2 days of EJS induction. Plotted are the mean ±SD. Statistical significance was determined by one-way ANOVA with Tukey’s multiple comparisons test;****p<0.0001. Numbers of wing discs analyzed for each time point are as follows (**Day**: n [EJS]; n [EJS + *Dronc*-i]): (**1**: 82; 78), (**2**: 35; 29), (**3**: 18; 7). **(C)** Confocal image of hemocytes labeled with *hml-*dsRed found on EJS tumors after 5 days of EJS induction (gray, magenta); notice the colocalization of the EdU incorporation (green) with the nuclear hemocyte marker (arrows; white indicates overlap) **(D and E)** Number of EdU^+^ hemocytes as a function of total number of hemocytes in control (EJS, blue) and EJS + UAS-*Dronc*-RNAi (EJS + *Dronc*-i, orange) after 1 (D) and 2 (E) days of EJS induction. Pearson R coefficients displayed. Statistical significance between the correlation coefficient difference was determined by a Fisher-Z-Transformation, via a two-tailed test. ns – not significant, p = 0.0569; *** p = 0.0077. Numbers of wing discs analyzed for each time point are as follows (**Day**: n [EJS]; n [EJS + *Dronc*-i]): (**1**: 18; 13), (**2**: 8; 8). **(F)** Representative confocal images of hemocytes labeled with *hml*-dsRed (gray) in control tumors (EJS), Dronc-deficient EJS tumors (EJS + *Dronc*-i), JNK-deficient EJS tumors (EJS + JNK^DN^), and both Dronc- and JNK-deficient EJS tumors (EJS + *Dronc*-i + JNK^DN^) after 1.5 days of EJS induction. Dashed white outline of wing discs obtained by tracing DAPI staining (not shown). Scale bar: 100μm. **(G)** Number of hemocytes per tumor in control tumors (EJS), Dronc-deficient EJS tumors (EJS + *Dronc*-i), JNK-deficient EJS tumors (EJS + JNK^DN^), and both Dronc- and JNK-deficient EJS tumors (EJS + *Dronc*-i + JNK^DN^) after 1.5 days of EJS induction. Full genotypes can be found in the MM. Statistical significances was determined by one-way ANOVA with Turkey’s multiple comparisons tests. ns – not significant p>0.05, **** p<0.0001. Numbers of wing discs analyzed for EJS tumors: 39; for EJS + *Dronc*-i tumors: 50; for EJS + JNK^DN^ tumors: 30; for EJS + *Dronc-*i + JNK^DN^: 29. **(H)** Number of hemocytes per tumor in control and Dronc-deficient EJS tumors in a genetic background with either wild-type hemocytes (EJS and EJS + *Dronc*-i) or hemocytes expressing the pro-apoptotic factor reaper (EJS + *hml>rpr* and EJS + *Dronc*-i + *hml>rpr*) after 1 day of EJS induction. Statistical significances was determined by one-way ANOVA with Turkey’s multiple comparisons tests. ns – not significant p>0.05, **** p<0.0001. Numbers of wing discs analyzed for EJS tumors: 21; for EJS + *hml*>*rpr* tumors: 21; for EJS + *Dronc*-i tumors: 25; for EJS *hml*>*rpr* + *Dronc*-i: 29. **(I)** Relative size of control tumors (EJS) and EJS tumors in a genetic background with hemocytes expressing the pro-apoptotic gene *reaper* (EJS + *hml*>*rpr*) after 3 days of EJS induction. Plotted are the mean ±SD of relative tumor sizes at each time point. Control tumors were used for normalization. Statistical significance was determined by using an unpaired Student’s t-test; **** p<0.0001. Numbers of wing discs analyzed for EJS tumors: 29; for EJS + *hml*>*rpr* tumors: 30. **(J)** Relative size of Dronc-deficient tumors (EJS + *Dronc*-i) and EJS + *Dronc*-i tumors in a genetic background with hemocytes expressing the pro-apoptotic gene *reaper* (EJS + *Dronc*-i + *hml*>*rpr*) after 3 days of EJS induction. Plotted are the mean ±SD of relative tumor sizes at each time point. EJS + *Dronc*-i tumors were used for normalization. Statistical significance was determined by using an unpaired Student’s t-test; **** p<0.0001. Numbers of wing discs analyzed for EJS + *Dronc*-i tumors: 30; for EJS + *Dronc*-i + *hml*>*rpr* tumors: 34.

Since many of the features of caspase-deficient EJS tumors were previously correlated with the upregulation of JNK signaling in transformed cells, we evaluated the contribution of this pathway to the excess of DTAMs. As previously described for other EJS features, the elimination of JNK activity in *Dronc*-deficient EJS tumors abolished the increased number of DTAMs (Figure 5F,G). These results showed that the excess of DTAMs in *Dronc*-deficient EJS tumors is mediated by the exacerbation of JNK signaling.

### The presence of functional tumor-associated macrophages contributes to tumor growth

Immune cells in the tumor microenvironment can either promote or inhibit tumor growth both in *Drosophila* and mammalian models (Cordero et al., 2010; Pastor-Pareja et al., 2008; Perez et al., 2017; Shalapour and Karin, 2015). To better define the role of DTAMs in the growth of EJS tumors, we attempted to eliminate them by expressing the pro-apoptotic gene *reaper*. This genetic manipulation successfully killed the hemocytes in wild-type larvae without tumors (Charroux and Royet, 2009; Pastor-Pareja et al., 2008; Shia et al., 2009) (Figures S4) but it failed to affect the number of DTAMs on EJS tumors (Figure 5H). However, *reaper* overexpression caused a significant reduction in EJS tumor size (Figure 5I). Furthermore, this tumor reduction was not suppressed in caspase-deficient conditions (Figure 5J). This leads to the important conclusion that DTAMs play a pro-tumorigenic role in EJS tumors, which in turn can be disabled by expressing the pro-apoptotic factor *reaper*. Furthermore, these *reaper*-dependent effects occur without inducing apoptosis or compromising the ability of hemocytes to either proliferate or be recruited to the tumors (see Discussion).

## Discussion

Whereas the role of caspases has been exhaustively characterized during apoptosis, it remains largely unexplored in non-apoptotic cellular scenarios. Our work has uncovered novel tumor-suppressor functions linked to caspases independent of apoptosis, which limit JNK signaling and the number of tumor-associated macrophages in our oncogenic model. Importantly, these novel non-apoptotic caspase functions are required in virtually all of the transformed cells for preventing the exacerbation of malignant features, such as excess proliferation or defective cell differentiation. In parallel, our results demonstrate that the simultaneous hyperactivation of JNK and JAK/STAT pathways confer a signaling profile to EJS tumors highly reminiscent to that of cells during regeneration, which permanently retains tumor cells in an undifferentiated and proliferative state. Therefore, we conclude that EJS tumors can be considered a *bona fide* paradigm of open wound-like tumors in *Drosophila*. Together, our findings improve our basic understanding of caspase biology in tumors and how these enzymes can suppress oncogenic transformation through mechanisms independent of apoptosis.

### Caspases can act as tumor-suppressors through molecular mechanisms independent of apoptosis

The tumor suppressing activity of caspases has primarily been linked to their ability to activate the apoptotic program, and therefore, cell death (Hanahan and Weinberg, 2000, 2011; Olsson and Zhivotovsky, 2011). However, recent evidence has shown that these enzymes play instrumental roles in the regulation of alternative cellular functions beyond apoptosis (e.g. intracellular signaling regulation) that can have a significant impact on tumor biology (Fogarty and Bergmann, 2017; Portela and Richardson, 2013; Xu et al., 2018). Our findings demonstrate that non-apoptotic caspase activity can strongly determine the signaling profile of transformed cells and basic properties of the tumor microenvironment. Importantly, these caspase-dependent effects limit tumor growth and facilitate cell fate commitment in EJS tumors; therefore, they can be considered tumor-suppressor activities of caspases independent of apoptosis. This conclusion broadens the repertoire of caspase-mediated functions that can compromise tumor progression.

### Tumors can be formed mainly by caspase-activating cells that do not die

Our caspase activation analyses in EJS tumors show that virtually all of the cells in a tumor can either undergo cycles of caspase activation/deactivation without completing the apoptosis program or alternatively, derive from caspase-activating cells that are not eliminated (Figure 1). Furthermore, subsequent functional genetic studies ascribe this caspase activation to the *Drosophila Caspase-2/9* ortholog known as *Dronc*, instead of the effector caspases. Supporting this conclusion, we establish that while *Dronc* deficiency potently aggravates the oncogenic features of EJS tumors, the lack of effector caspase activity fails to alter tumor properties. These findings indicate that novel tumor-suppressor activities linked to initiator caspases can be highly represented within tumors and can limit the malignant potential of transformed cells (Krelin et al., 2008; Puccini et al., 2013). Intriguingly, although the completion of apoptosis through effector caspases is irrelevant in our tumor setting, the activation of Dronc still relies on the apoptotic gene Apaf-1 (Dark in flies) (Figure 2F). This suggests that the basal assembly of the apoptosome is responsible to a large extent for the non-apoptotic caspase functions in our tumor model. Interestingly, recent studies have started to attribute similar non-apoptotic functions to Apaf-1 during the differentiation of muscle progenitor cells (H. Dehkordi et al., 2020).

### Non-apoptotic caspase activation compromises the growth and the malignant transformation of “open-wound” like tumors by limiting JNK signaling

The similarities between the cellular properties found in tumors and cells taking part in the regenerative process after injury suggested the hypothesis that tumors could behave like open wounds that never heal (Byun and Gardner, 2013; Dvorak, 1986, 2015; Ribatti and Tamma, 2018). During tissue regeneration and wound healing, the *Drosophila* cells forming the blastema display a stereotypical signaling profile that includes the simultaneous activation of JAK-STAT and JNK pathways (Ahmed-de-Prado et al., 2018; Santabarbara-Ruiz et al., 2015; Worley et al., 2018). The transient and cooperative activity of both signaling pathways elicits cell re-specification in damaged tissues (Ahmed-de-Prado et al., 2018; Worley et al., 2018). Our data indicate that the upregulation of JAK/STAT and EGFR pathways in EJS tumors permanently activates JNK signaling, which in turn eliminates the expression of cell identity markers and causes excess proliferation (Figure 4). Supporting this conclusion, deprivation of JNK signaling can largely rescue most of the malignant features (patterning defects, tissue hyperplasia, and DTAM recruitment) in EJS tumors. In this context, we also established that non-apoptotic caspase activation limits JNK signaling, thus preventing exacerbation of malignant properties. Considering the strong parallels between the *Drosophila* regeneration process and EJS tumor properties, we suggest that EJS tumors behave like open wounds that never heal. Importantly, the oncogenic potential of the JAK/STAT and EGFR cooperation has also been demonstrated in mammalian systems (Andl et al., 2004; Quesnelle et al., 2007; Wen et al., 2015).

### The interplay between caspases and JNK pathway is complex and highly context-dependent

Our experiments now provide seminal evidence *in vivo* indicating that initiator caspases can limit JNK signaling and the malignant exacerbation of oncogenic features. Although this is the first time that the non-apoptotic ability of caspases has been shown to moderate JNK signaling in a tumor model, the cleavage of the JNK Interacting Protein-1 (JIP1) mediated by effector caspases ameliorates JNK signaling in mammalian cells during apoptosis (Vaishnav et al., 2011). Additionally, a recent report has also shown that caspases also limit a major stress response MAPK pathway similar to JNK to ensure developmental progression and growth in *C. elegans* (Weaver et al., 2020). These observations open the intriguing possibility that specific substrates of Dronc within the JNK pathway only become available in EJS tumors. In line with this possibility, JNK activation can be achieved differently in distinct tumor models: whereas EJS tumors show JNK activation that is membrane receptor and ROS independent (Figure S3), JNK activation in *scrib*^−/−^/*Ras*^*V12*^ tumors fully relies on these factors (Fogarty et al., 2016; Perez et al., 2017). Beyond the specific relationship between caspases and JNK signaling, there are alternative causes that could facilitate the novel tumor suppressing activity of caspases in EJS tumors. One of these could be the strength of the caspase activation. Supporting this hypothesis, it has been shown that JAK/STAT activation can promote cell survival by inducing the expression of the caspase inhibitor Diap-1 (Betz et al., 2008; Recasens-Alvarez et al., 2017), and therefore could maintain caspase activation under sublethal thresholds in EJS tumors. Although further studies outside of the scope of this manuscript are needed to fully understand the complex interplay between caspases, JNK signaling, and tumor progression, our data provide novel molecular leads to understanding the non-apoptotic regulation and highly context-dependent function of caspases in tumors.

### The non-apoptotic activity of initiator caspases modulates the recruitment and proliferation of tumor-associated macrophages by limiting JNK signaling

Although the influence of caspase activation on the tumor microenvironment has been acknowledged across different animal species (Legrand et al., 2019), it is not fully understood. Recently, it has been shown that caspase activation in a *Drosophila* model of *scrib*^−/−^/*Ras*^*V12*^ tumors leads to ROS generation, which subsequently attracts the *Drosophila* macrophage-like cells (hemocytes) towards the transformed cells. Once the *Drosophila* tumor-associated macrophages (DTAMs) become active *in situ*, they can release Eiger (TNF-α), thus promoting JNK activation and tumor growth (Fogarty et al., 2016; Perez et al., 2017). In striking contrast, in EJS tumors, non-apoptotic caspase activation limits the number of DTAMs and tumor growth (Figure 5). Furthermore, these disparities appear despite the fact that the production of ROS in EJS tumors is linked to caspases. Beyond these context-dependent differences, our results show that the excess of DTAMs in EJS tumors without caspase activity largely relies on the hyperactivation of JNK signaling. Supporting this conclusion, deficiency in JNK signaling significantly reduced the number of DTAMs in caspase-deficient EJS tumors (Figure 5G). A plausible molecular explanation for these JNK effects could be related to the upregulation of Upd ligands (orthologs of mammalian interleukins) downstream of this pathway (Figure 4), since it has been shown that the release of these soluble factors can stimulate the recruitment and cell proliferation of immune cells (O’Shea and Plenge, 2012; Pastor-Pareja et al., 2008) (Figure 6). In line with this hypothesis, the total number of DTAMs in caspase-deficient EJS tumors is strongly correlated with the number of DTAMs positively labelled with the S-phase marker EdU (Figure 5E). This is an important observation, since it indicates for the first time that the excess of DTAMs in caspase-deficient EJS tumors can be linked, at least to some extent, to the *in situ* proliferation of the immune cells on the tumor. Together, previous literature (Pastor-Pareja et al., 2008; Perez et al., 2017) and our data demonstrate the ability of caspase activation to modulate the cellular configuration of the tumor microenvironment through multiple molecular mechanisms.

**Figure 6.**
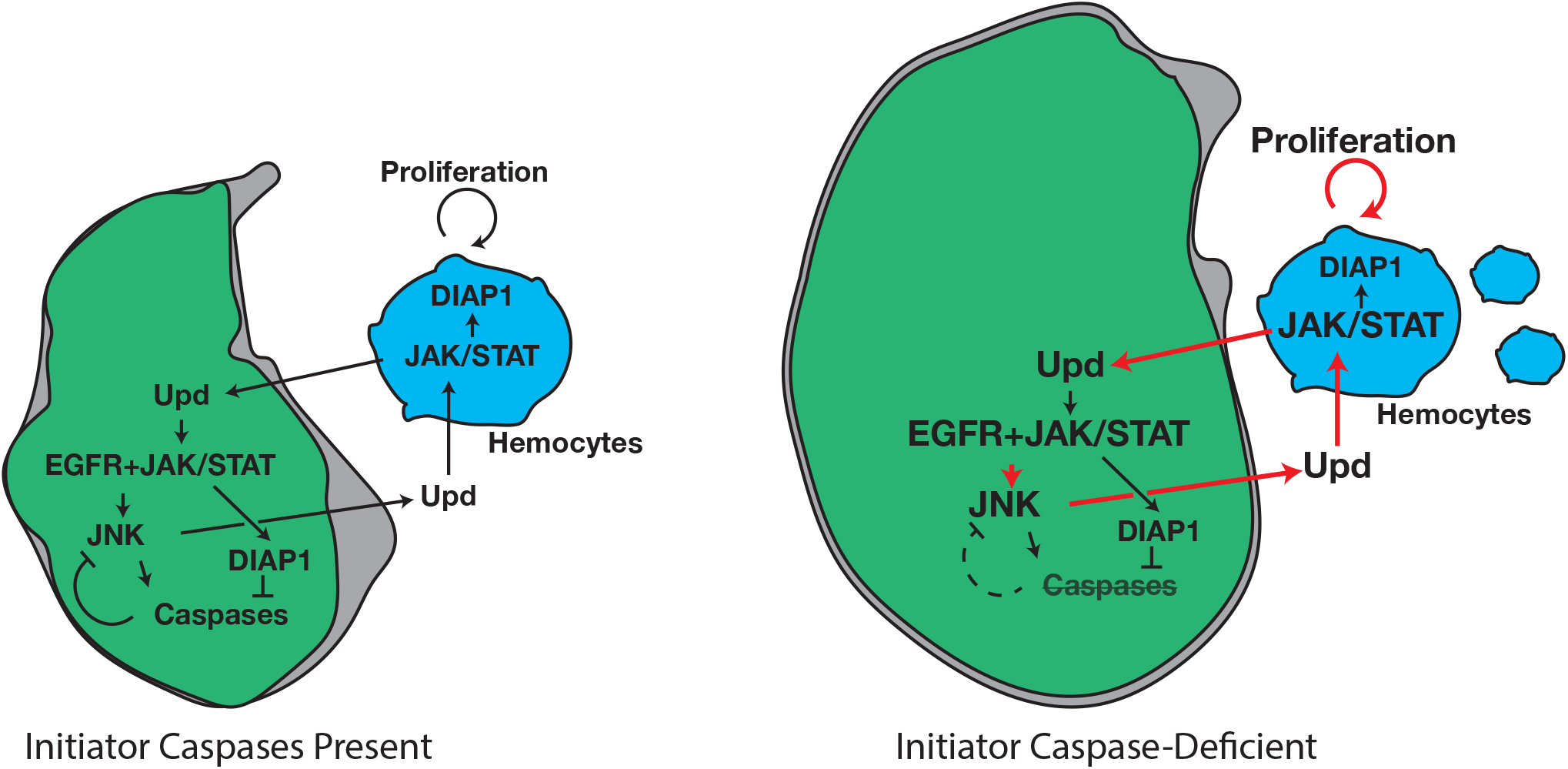
Initiator caspases can act as tumor suppressors in an open wound-like EJS tumor by negatively regulating the JNK pathway. Diagrams indicating the signaling profile, growth (size), and hemocyte interactions in EJS (left) and caspase-deficient EJS tumors (right). Notice the positive feedback loop reinforced by the absence of caspase activity (thicker red arrows) that promotes JNK signaling, hemocyte proliferation, and tumor growth.

### Functional immune cells on tumors sustains oncogenic growth

The presence of immune cells in the tumor microenvironment is at the forefront of the defense against tumors; however, there are also examples in which immune cells can fuel tumor growth, mainly through the release of soluble molecules that either act as trophic factors and/or protect tumor cells against cell death mechanisms (Cordero et al., 2010; Gonzalez et al., 2018; Pastor-Pareja et al., 2008; Perez et al., 2017). Our data suggest that the excess of DTAMs can contribute to the growth of EJS tumors, but only if they are functionally active. Supporting this conclusion, overexpression of the pro-apoptotic factor Reaper in hemocytes markedly compromised tumor growth but failed to compromise DTAM viability in our tumor scenario (Figure 5I,J). This striking result has important implications. First, *reaper*-expressing DTAMs must still be receive proliferative signals from the EJS-transformed cells, since their number does not decrease after expressing *reaper*. Second, the DTAMs must receive survival cues from their interaction with the tumor cells, since like in wild-type conditions, they would otherwise die due to *reaper* expression. Third, although the survival and proliferative potential of *reaper*-expressing DTAMs is not compromised, their tumor-promoting ability is severely affected since EJS tumors are reduced in size, and therefore *reaper* overexpression disables DTAMs ability to sustain malignant tumor growth.

At the molecular level, our observations are compatible with the bidirectional secretion of Upd ligands from tumor cells and DTAMs, and the subsequent activation of the JAK/STAT pathway in both cell types. Whereas the release of Upd ligands from the EJS transformed cells (Figure 6) could activate and induce the proliferation of recruited DTAMs (Pastor-Pareja et al., 2008; Woodcock et al., 2015; Yang et al., 2015), the secretion of the same soluble ligands from DTAMs could fuel tumor growth (Amoyel et al., 2014). In parallel to the proliferation effects, the activation of the JAK/STAT pathway could provide survival cues to both DTAMs and tumor cells since this signaling cascade can induce the expression of apoptosis inhibitors (Betz et al., 2008; Recasens-Alvarez et al., 2017). Based on this, we propose that the mutual activation of the JAK/STAT pathway in EJS transformed cells and DTAMs could establish a positive feedback loop that ultimately facilitates tumor progression (Figure 6). Importantly, this feedback loop would be negatively regulated by the caspase-dependent tuning of JNK signaling. Considering that the cross-talk between TAMs and transformed cells, as well as the pro-proliferative and pro-survival activities of EGFR and JAK/STAT pathways are represented in mammalian systems (Fisher et al., 2014), our *Drosophila* findings could be potentially relevant to understand the function of caspases in comparable human tumors.

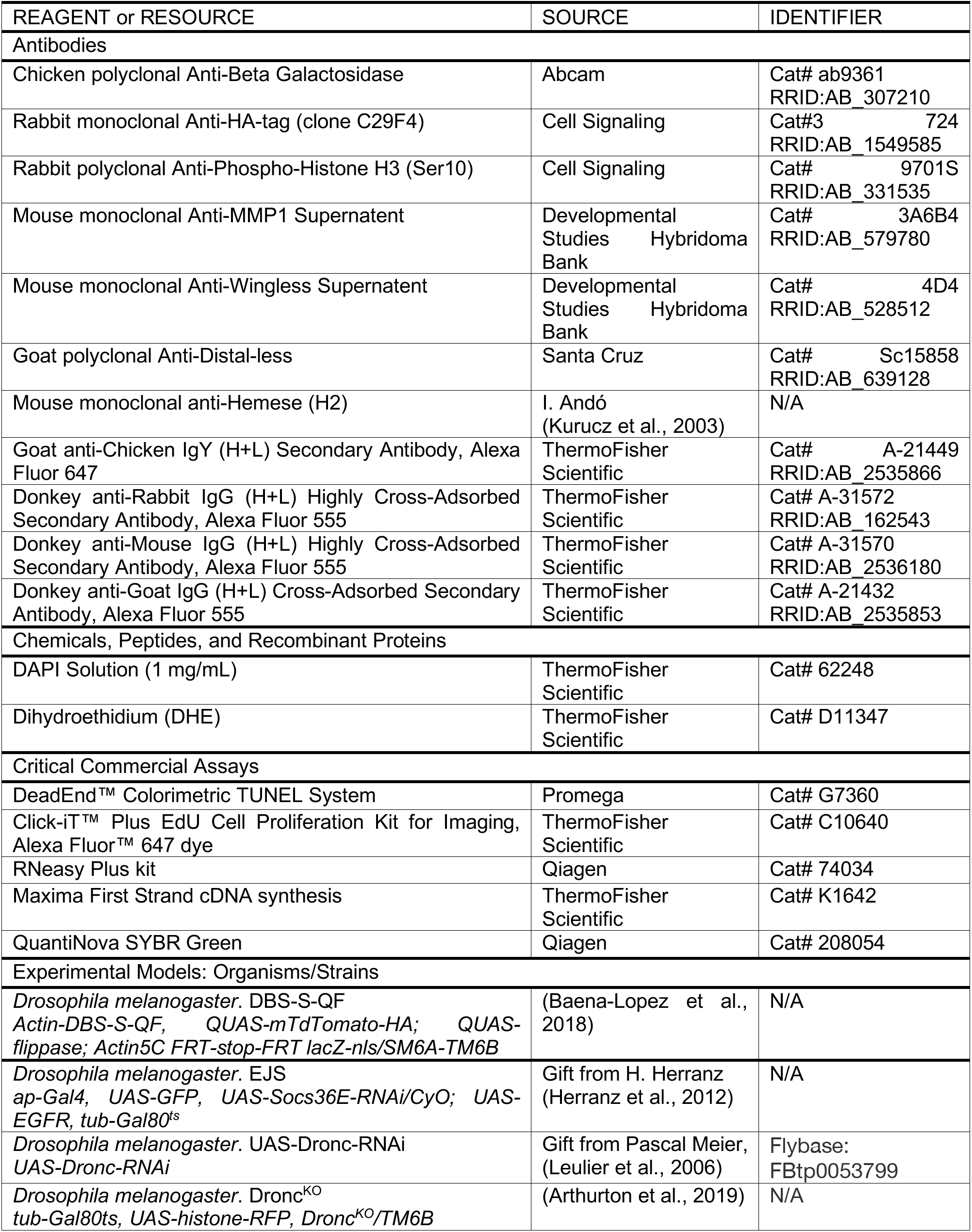
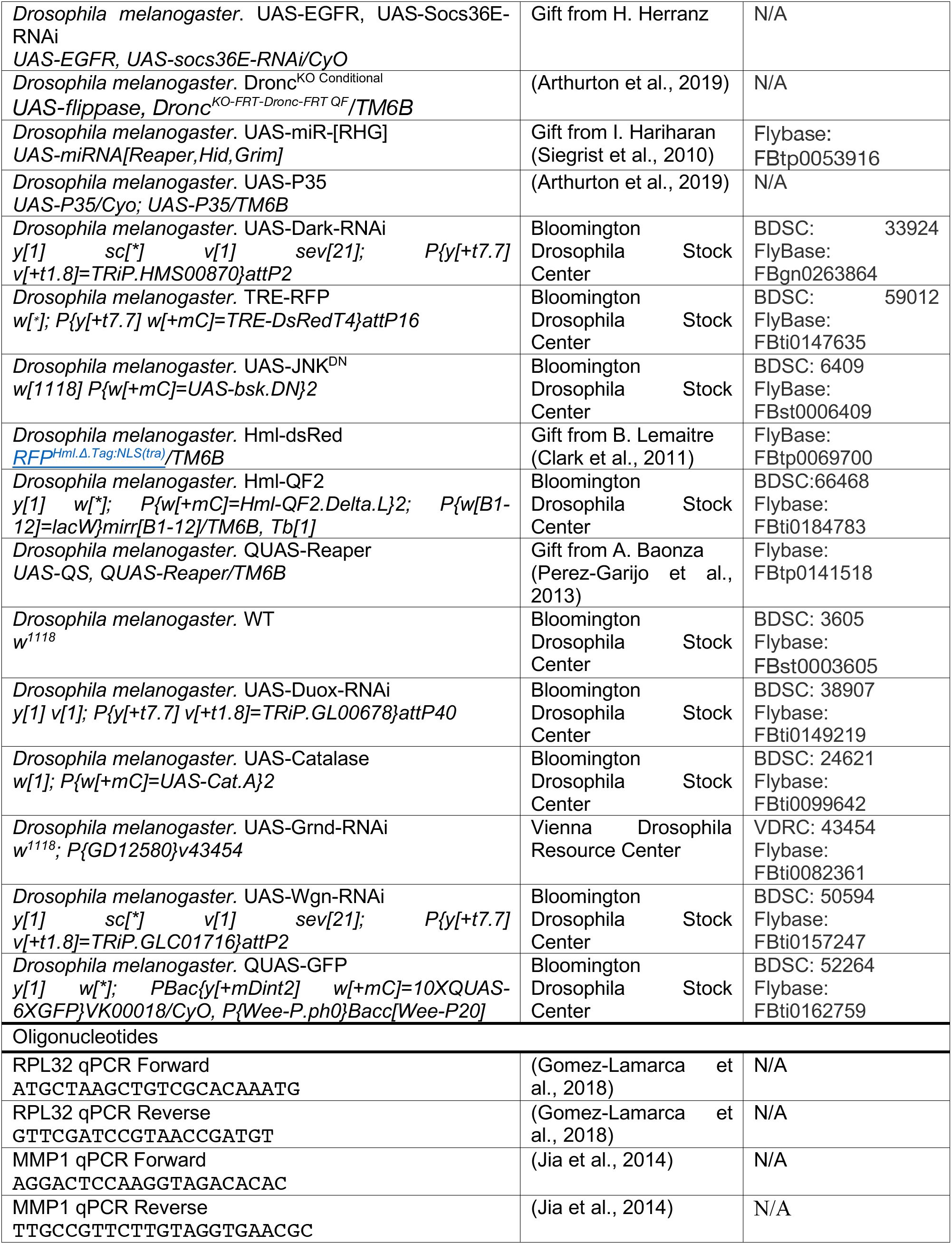
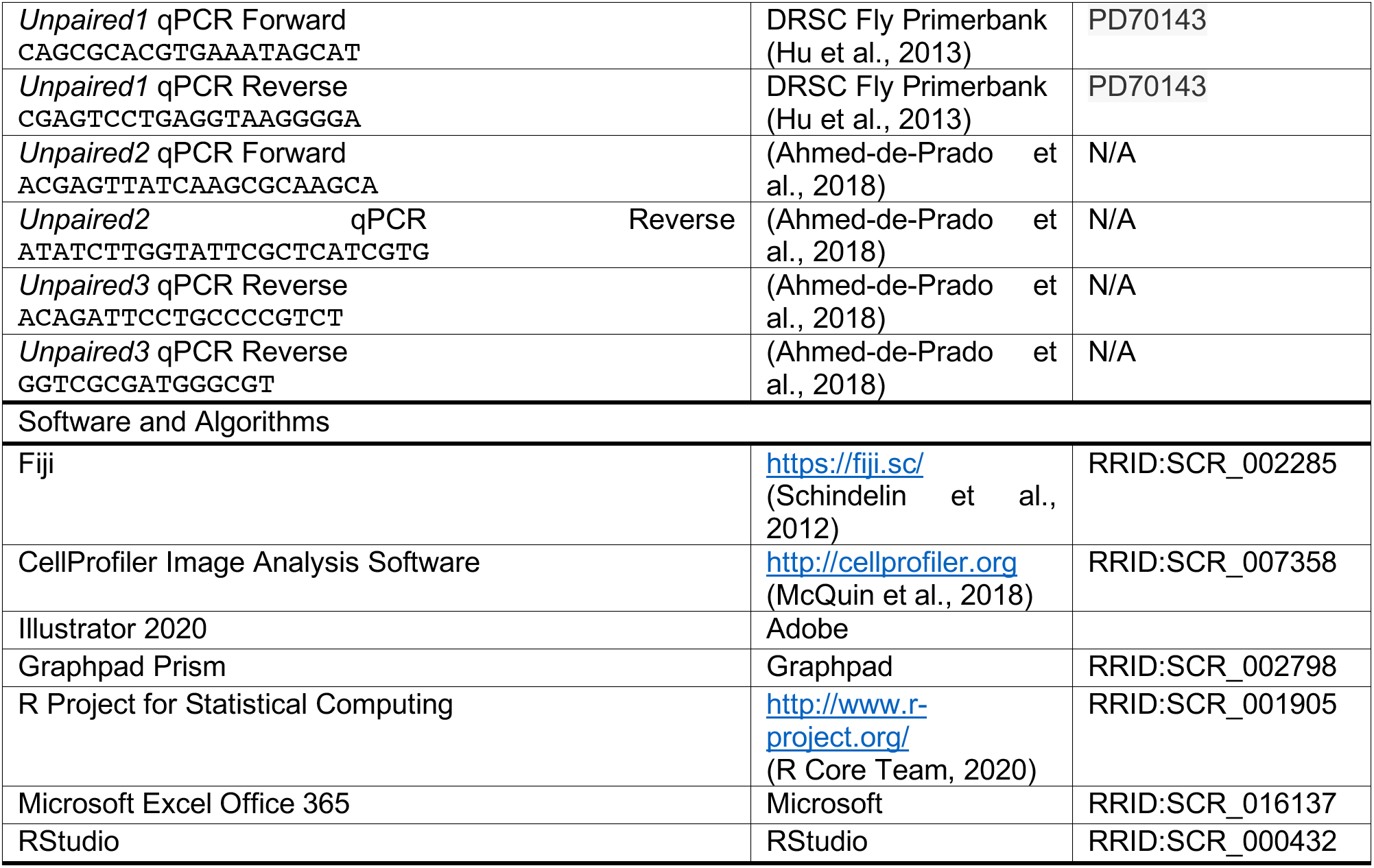
KEY RESOURCES TABLE.

## Resource Availability

### Contact for Reagent and Resource Sharing

Key fly strains will be deposited in Bloomington’s public repository soon after publication, but they will also be distributed upon request. Further information and requests for resources and reagents should be directed to and will be fulfilled by the Lead Contact, Luis Alberto Baena-Lopez

## Experimental Model and Subject Details

### *Drosophila* stocks and husbandry

All *Drosophila* stocks were maintained at 20°C or 18°C in *Drosophila* culture medium (0.77% agar, 6.9% maize, 0.8% soya, 1.4% yeast, 6.9% malt, 1.9% molasses, 0.5% propionic acid, 0.03% ortho-phosphoric acid, and 0.3% nipagin) was used to maintain flies in vials.

## Method Details

### Tumor induction

For experiments involving induction of tumor formation during larval stages, Gal80^ts^ was used to temporally control overexpression of oncogenes. Flies were reared and crossed at 18°C, inhibiting Gal4 activity. 30–40 virgins were crossed to 10–15 males for each genotype and experiment. Crosses were flipped twice a day (morning and evening) into fresh vials of food, to ensure larvae used in experiments were of similar age. One day before larvae entered the wandering third instar stage (9 full days in our fly incubators and experimental environment), larvae were shifted to 29°C for up to 5 days, depending upon the experiment.

### Immunohistochemistry

Experimental specimens dissected were larvae crawling along the walls of the vial. Wing discs were dissected according to standard protocols in ice-cold PBS and collected in 4% formaldehyde in 1x PBS on ice to prevent potential hemocyte dissociation from the tissue. Fixation occurred for an additional 20 minutes after dissections at room temperature. Discs were washed in PBS-TX (1x PBS, 0.3% Triton X-100) and blocked in blocking solution (3% BSA and 0.5% sodium azide in PBS-TX).

Discs were incubated while shaking overnight at 4°C with primary antibodies in blocking solution. The following primary antibodies and concentrations were used in this study: chicken anti-Beta-galactosidase (1:500, Abcam, RRID:AB_307210), rabbit anti-HA-tag (1:1000, Cell Signaling, RRID:AB_1549585), rabbit anti-Phospho-histone H3 (1:100, Cell Signaling, RRID:AB_331535), mouse anti-MMP1 (1:200, Developmental Studies Hybridoma Bank, RRID:AB_579780), mouse anti-Wingless (1:200), Developmental Studies Hybridoma Bank, RRID:AB_528512), goat anti-Distal-less (1:100, Santa Cruz, RRID:AB_639128), and mouse anti-Hemese (H2) (1:500, gift from I. Andó).

After washing, discs were incubated with conjugated secondary anti-bodies for 2 hours at room temperature. The following secondary antibodies were used in this study, all from ThermoFisher Scientific and used at a concentration of (1:200): Goat anti-chicken Alexa Fluor 647 conjugated, donkey anti-rabbit, -mouse, and -goat Alexa Fluor 555 conjugated. DAPI (1:1000) was added to a 15-minute washing step after secondary antibody incubation. After washing, discs were rinsed with 1x PBS, and then incubated in a 50% glycerol solution in 1x PBS for 1 hour. Discs were then incubated in an 80% glycerol solution in PBS for at least one hour. Discs were removed from the carcasses and mounted in 80% glycerol. Control and experimental samples were mounted on the same slide to control for sample compression. Samples were then covered with a 1.5H coverslip, secured with nail polish, and imaged immediately or stored at 4°C.

### DHE labeling

DHE labeling of ROS was conducted according to a previously described protocol (Owusu-Ansah et al., 2008), including the optional fixation in 4% formaldehyde in 1x PBS.

### TUNEL staining

As for immunohistochemistry, wing discs were dissected in ice-cold PBS and fixed in 4% formaldehyde in PBS for 20 minutes. *In situ* detection of fragmented genomic DNA was then performed according to methods established previously (Galasso et al., 2020), using the DeadEnd Colorimetric TUNEL (Terminal transferase-mediated dUTP nick-end labeling) system (Promega).

### EdU staining

For EdU staining, larvae were selected and dissected as described above (please see immunohistochemistry section above). Instead of fixative, discs were collected in ice cold PBS. Dissection times were kept under 20 minutes. After dissection, the PBS was replaced with a 0.1mg/mL solution of EdU in PBS for a 5-minute incubation. After incubation, discs were rinsed with PBS and then fixed in 4% formaldehyde in 1x PBS for 20 minutes. Subsequent steps followed instructions according to the manufacturer (ThermoFisher, Click-iT Edu Kit).

### Fluorescence microscopy and image analysis

Images were acquired using an inverted FV1200 Olympus microscope, using 10x air, 20x air, or 60x oil objective lenses, depending on the experiment. Unless otherwise indicated, the entire wing disc was imaged in all three dimensions, using the optimal step size determined by the Olympus Fluoview software. For images where relative intensities were being measured, a constant laser power was used throughout genotypes and experiments. Post-acquisition imaging processing was performed using Fiji (Schindelin et al., 2012) and CellProfiler image analysis software (CellProfiler).

To measure EJS tumor sizes, maximum intensity projections of each z-stack were generated using Fiji to produce 2D images. Tumor sizes were measured using CellProfiler’s IdentifyPrimaryObjects module.

To measure nuclei sizes and density in EJS tumors, the midpoint of a disc was determined in the z-axis, and a 60x image of the DAPI channel was acquired in a similar location amongst all discs, in the ventral compartment. Using CellProfiler, nuclei sizes were segmented and measured using the IdentifyPrimaryObjects module. To measure nuclei density, images were shuffled and blinded to the investigator for quantification. A 100μm^2^ area was selected in the center of all images and the number of nuclei were counted manually using Fiji.

To quantify the numbers of hemocytes present on EJS tumors, maximum intensity projections of each z-stack were generated using Fiji to produce 2D images. After a tumor was identified and outlined using CellProfiler, a mask was applied on the channel containing hemocyte nuclei (visualized using hml-dsRed) to restrict counting to hemocytes present only on the wing disc. Individual hemocyte nuclei were segmented, identified, and counted using CellProfiler’s IdentifyPrimaryObjects module.

To quantify proliferation in EJS tumors, maximum intensity projections of each z-stack were generated using Fiji to produce 2D images. For PH3 based measurements, after a tumor was identified and outlined using CellProfiler, a mask was applied on the channel containing PH3 staining to restrict counting to cells in the EJS tumor. Individual hemocyte nuclei were segmented, identified, and counted using CellProfiler’s IdentifyPrimaryObjects module. For EdU based measurements, thresholding in Fiji was used to determine both EJS tumor area and EdU^+^ area.

To measure the intensity of *Tre-RFP* and antibody staining for MMP1 in EJS tumors, maximum intensity projections of each z-stack were generated using Fiji to produce a 2D image. For MMP1 stained samples, mean intensities in EJS tumors were measured using Fiji. For *Tre-RFP* samples, individual nonzero pixel values were extracted from each image using the Save XY Coordinates function and imported into R Project (R Core Team, 2020) using RStudio. Pixel intensity values below a certain background value (determined using the Threshold function in Fiji) were removed from the data set. A density curve of pixel intensities was calculated for each disc, along with the mode intensity using the ggridges package (Wilke, 2020). The standard deviation of the modes for each genotype was also calculated using R.

### Real-time quantitative PCR

Larvae were collected and identified in the same manner as for immunostaining after 3 days of tumor induction. Larvae were dissected in one well of a 9-well dissection plate on ice, and inverted carcasses were collected in a separate well containing PBS. Once 15–20 larvae were collected, wing discs were carefully separated from the carcass, disposing of the carcass. Dissection time was limited to 20–30 minutes. Cleaned discs were then transferred to a clean 0.5mL Eppendorf tube, using a P20 micropipette tip. Cut the tip off the P20 micropipette tip and coat the tip by pipetting the contents of the remaining carcasses up and down several times to prevent discs from sticking to the tip. After the discs were transferred, the PBS was replaced with 100μl of RNA lysis buffer. Tissue was lysed by short bursts of vortexing using a tabletop vortex mixer. Samples of similar genotypes can be frozen at this point in liquid nitrogen and pooled together if necessary. RNA was subsequently extracted using an RNA Easy Plus kit following the manufacturer’s instructions (Qiagen, 74034). 500ng of total RNA of each sample was then used for reverse transcription, according to manufacturer instructions (Thermo Fischer Scientific Maxima First Strand cDNA Synthesis Kit – K1671). Q-PCR was then performed using a QuantiNova SYBER Green PCR Kit (Qiagen, Cat# 208054) and a Rotor-Gene Q Rea-time PCR cycler (Qiagen). Data was analyzed using the comparative CT method (Livak and Schmittgen, 2001) with *RPL32* used as a housekeeping gene for internal control. Primers used for qPCR can be found in the STAR Key Resources Table.

### Quantification and statistical analysis

All data was processed and analyzed using R, Graphpad Prism 8, and Microsoft Excel. Unpaired Student’s t-tests, or one-way ANOVA with Tukey’s multiple comparisons test were used to compare values between genotypes for univariate data. To compare Pearson correlation coefficients for multivariable data, a Fisher-Z-Transformation two-tailed test was used (Diedenhofen and Musch, 2015).

### Figure Generation

Figures were generated using Adobe Illustrator 2020. For confocal images, wing discs were arranged to be in the same orientation such that the anterior direction is to the left and the dorsal side is to the top of the page. When confocal images were rotated, a dark rectangular background was added to create regularly shaped figures.

### Information of detailed genotypes

The detailed genotypes of each experiment are described below:

**Figure 1:**

**(B)** *Actin-DBS-S-QF, QUAS-mTdTomato-HA*; *QUAS-flippase*/+; *Actin5C FRT-stop-FRT lacZ-nls*/+

**(D and E)** *Actin-DBS-S-QF* (Baena-Lopez et al., 2018)*, QUAS-mTdTomato-HA* (BL30118); *ap-Gal4, UAS-GFP, UAS-Socs36E-RNAi* (gift from H. Herranz, Herranz et al., 2012)/*QUAS-flippase* (BL30126)*; UAS-EGFR, tub-Gal80*^*ts*^ (gift from H. Herranz, Herranz et al., 2012)*/Actin5C FRT-stop-FRT lacZ-nls* (BL6355)

**Figure 2:**

**(A, B, and C)** EJS: *ap-Gal4, UAS-GFP, UAS-Socs36E-RNAi*/+; *UAS-EGFR, tub-Gal80*^*ts*^/+

EJS + *Dronc-i: ap-Gal4, UAS-GFP, UAS-Socs36E-RNAi/UAS-Dronc-RNAi* (gift from P. Meier, Leulier et al., 2006)*; UAS-EGFR, tub-Gal80*^*ts*^/+

**(D)** *Dronc*^+/−^: *ap-Gal4/UAS-EGFR, UAS-Socs36E-RNAi* (gift from H. Herranz)*; tub-Gal80*^*ts*^ (BL7019)*, UAS-histone-RFP* (BL56555)*, Dronc*^*KO*^ (Arthurton et al., 2019)/+

*Dronc*^−/−^: *ap-Gal4/UAS-EGFR, UAS-Socs36E-RNAi; tub-Gal80*^*ts*^,*UAS-histone-RFP, Dronc*^*KO*^/*UAS-flippase* (BL8209)*, Dronc*^*KO-FRT-Dronc-FRT QF*^ (Arthurton et al., 2019)

**(E)** EJS: *ap-Gal4, UAS-GFP, UAS-Socs36E-RNAi*/+; *UAS-EGFR, tub-Gal80*^*ts*^/+

EJS + *Dronc-i: ap-Gal4, UAS-GFP, UAS-Socs36E-RNA*/*Dronc-RNAi; UAS-EGFR, tub-Gal80*^*ts*^/+

*EJS* + *miR-[RHG]: ap-Gal4, UAS-GFP, UAS-Socs36E-RNAi/UAS-miRNA[Reaper,Hid,Grim]* (gift from I. Hariharan, Siegrist et al., 2010)*; UAS-EGFR, tub-Gal80*^*ts*^/+

EJS + P35: *ap-Gal4, UAS-GFP, UAS-Socs36E-RNAi/UAS-P35* (BL5072)*; UAS-EGFR, tub-Gal80*^*ts*^/*UAS-P35* (BL5073)

**(F)** EJS: *ap-Gal4, UAS-GFP, UAS-Socs36E-RNAi*/+; *UAS-EGFR, tub-Gal80*^*ts*^/+

EJS + *miR-[RHG]: ap-Gal4, UAS-GFP, UAS-Socs36E-RNAi/UAS-miRNA[Reaper,Hid,Grim]; UAS-EGFR, tub-Gal80*^*ts*^/+

EJS + P35: *ap-Gal4, UAS-GFP, UAS-Socs36E-RNAi/UAS-P35; UAS-EGFR, tub-Gal80*^*ts*^/UAS-P35

EJS + Dark-i: *ap-Gal4, UAS-GFP, UAS-Socs36E-RNAi*/+; *UAS-EGFR, tub-Gal80*^*ts*^/*UAS-Dark-RNAi* (BL33924)

**Figure 3:**

**(A, B, C, D, and E)** EJS: *ap-Gal4, UAS-GFP, UAS-Socs36E-RNAi*/+; *UAS-EGFR, tub-Gal80*^*ts*^/+

*EJS* + *Dronc-i: ap-Gal4, UAS-GFP, UAS-Socs36E-RNAi/Dronc-RNAi; UAS-EGFR, tub-Gal80*^*ts*^/+

**Figure 4:**

**(A and B)** EJS: *ap-Gal4, UAS-GFP, UAS-Socs36E-RNAi*/+; *UAS-EGFR, tub-Gal80*^*ts*^/+

*EJS* + *Dronc-i: ap-Gal4, UAS-GFP, UAS-Socs36E-RNAi/Dronc-RNAi; UAS-EGFR, tub-Gal80*^*ts*^/+

**(C and D)** EJS: *ap-Gal4, UAS-GFP, UAS-Socs36E-RNAi/Tre-RFP* (BL59012)*; UAS-EGFR, tub-Gal80*^*ts*^/+

*EJS* + *Dronc-i: ap-Gal4, UAS-GFP, UAS-Socs36E-RNAi/Dronc-RNAi, Tre-RFP; UAS-EGFR, tub-Gal80*^*ts*^/+

**(E)** EJS: *ap-Gal4, UAS-GFP, UAS-Socs36E-RNAi*/+; *UAS-EGFR, tub-Gal80*^*ts*^/+

EJS + *Dronc*-i: *ap-Gal4, UAS-GFP, UAS-Socs36E-RNAi/Dronc-RNAi; UAS-EGFR, tub-Gal80*^*ts*^/+

**(F and H)** WT: *w*^*1118*^ (BL3605)

EJS: *ap-Gal4, UAS-GFP, UAS-Socs36E-RNAi*/+; *UAS-EGFR, tub-Gal80*^*ts*^/+

EJS + *Dronc*-i*: ap-Gal4, UAS-GFP, UAS-Socs36E-RNAi/Dronc-RNAi; UAS-EGFR, tub-Gal80*^*ts*^/+

EJS + JNK^DN^*: UAS-bsk*^*DN*^ (BL6409)*; ap-Gal4, UAS-GFP, UAS-Socs36E-RNAi*/+; *UAS-EGFR, tub-Gal80*^*ts*^/+

EJS + JNK^DN^ + *Dronc*-i*: UAS-bsk*^*DN*^; *ap-Gal4, UAS-GFP, UAS-Socs36E-RNAi/Dronc-RNAi; UAS-EGFR, tub-Gal80*^*ts*^/+

**(J and K)** WT: *w*^*1118*^

EJS: *ap-Gal4, UAS-GFP, UAS-Socs36E-RNAi/+; UAS-EGFR, tub-Gal80*^*ts*^/+

*EJS* + *Dronc-i: ap-Gal4, UAS-GFP, UAS-Socs36E-RNAi/Dronc-RNAi; UAS-EGFR, tub-Gal80*^*ts*^/+

*EJS* + *JNK*^*DN*^: *UAS-bsk*^*DN*^; *ap-Gal4, UAS-GFP, UAS-Socs36E-RNAi*/+; *UAS-EGFR, tub-Gal80*^*ts*^/+

**Figure 5:**

**(A, B, D, and E)** EJS: *ap-Gal4, UAS-GFP, UAS-Socs36E-RNAi*/+; *UAS-EGFR, tub-Gal80*^*ts*^/*hml-dsRed*^*nls*^ (gift from B. Lemaitre, Clark et al., 2011)

*EJS* + *Dronc-i: ap-Gal4, UAS-GFP, UAS-Socs36E-RNAi/Dronc-RNAi; UAS-EGFR, tub-Gal80*^*ts*^/*hml-dsRed*^*nls*^

**(C)** *ap-Gal4, UAS-GFP, UAS-sSocs36E-RNAi/+; UAS-EGFR, tub-Gal80*^*ts*^/*hml-dsRed*^*nls*^

**(F and G)** EJS: *ap-Gal4, UAS-GFP, UAS-Socs36E-RNAi*/+; *UAS-EGFR, tub-Gal80*^*ts*^/*hml-dsRed*^*nls*^

*EJS* + *Dronc-i: ap-Gal4, UAS-GFP, UAS-Socs36E-RNAi/Dronc-RNAi; UAS-EGFR, tub-Gal80*^*ts*^/*hml-dsRed*^*nls*^

*EJS* + *JNK*^*DN*^: *UAS-bsk*^*DN*^; *ap-Gal4, UAS-GFP, UAS-Socs36E-RNAi*/+; *UAS-EGFR, tub-Gal80*^*ts*^/*hml-dsRed*^*nls*^

*EJS* + *JNK*^*DN*^ + *Dronc-i: UAS-bsk*^*DN*^; *ap-Gal4, UAS-GFP, UAS-Socs36E-RNAi/Dronc-RNAi; UAS-EGFR, tub-Gal80*^*ts*^/*hml-dsRed*^*nls*^

(H, I and J)

EJS: *ap-Gal4, UAS-GFP, UAS-Socs36E-RNAi*/+; *UAS-EGFR, tub-Gal80*^*ts*^/+

EJS + hml>rpr: *ap-Gal4, UAS-GFP, UAS-Socs36E-RNAi/hml-QF2* (BL66468)*; UAS-EGFR, tub-Gal80*^*ts*^/*UAS-QS, QUAS-Reaper* (gift from A. Baonza, Perez-Garijo et al., 2013)

*EJS* + *Dronc-i: ap-Gal4, UAS-GFP, UAS-Socs36E-RNAi/Dronc-RNAi; UAS-EGFR, tub-Gal80*^*ts*^/+

EJS + hml>rpr + *Dronc*-i: *ap-Gal4, UAS-GFP, UAS-socs36E-RNAi/hml-QF2, Dronc-RNAi; UAS-EGFR, tub-Gal80*^*ts*^/*UAS-QS, QUAS-Reaper*

## Author Contribution

D.C.X. designed, performed experiments and wrote the initial draft of the manuscript. L.A.B-L. initially conceptualized the project, designed experiments, secured funding, and reviewed and edited the manuscript. K.M.Y. secured funding, reviewed and edited the manuscript, and provided expertise and feedback.

## Acknowledgments

We would like to thank the following for flies and reagents: Hector Herranz (EJS flies and valuable input at the initial stages of the project); Pascal Meier (*UAS-Dronc-RNAi*); Iswar Hariharan (UAS-miR[RHG]); Bruno Lemaitre (Hml-dsRed); Antonio Baonza (QUAS-Reaper); Sonia Muliyil (UAS-Grindelwald-RNAi); István Andó (H2 antibody); the Bloomington *Drosophila* Stock Center and Vienna *Drosophila* Resource Center for fly strains. This work has been supported by Cancer Research UK C49979/A17516, the John Fell Fund from the University of Oxford (162/001) and NIH, NIDCR Intramural project ZIA DE000719. D.C.X. is a Ph.D student supported by the NIH NIDCR Intramural Research Program as part of the NIH-Oxford/Cambridge Scholars Program. is an NIH Distinguished Investigator and a Section Chief at NIDCR, NIH. L.A.B-L. is an Associate Professor at the University of Oxford, a CRUK Career Development Fellow and an Oriel College Hayward Fellow. All the authors declare no competing interest.

