## Supplementary information for "Widespread non-apoptotic activation of *Drosophila* Caspase-2/9 limits JNK signaling, macrophage proliferation, and growth of wound-like tumors"

**SUPPLEMENTARY FIGURE LEGENDS**

**Supplementary Figure 1.** **Density of nuclei in EJS + Dronc-i tumors is lower than control EJS tumors.**

The density of nuclei in control (EJS) and EJS + UAS-*Dronc*-RNAi (EJS + *Dronc*-i) tumors after 3 days of EJS induction, as measured by the number of nuclei per μm^2^. Plotted also are the mean ±SD. Statistical analysis performed by Student’s t-test. **** p<0.0001. Numbers of wing discs analyzed for EJS tumors: 34; for EJS + Dronc-i tumors: 38.

**Supplementary Figure 2. Inhibition of JNK signaling in EJS tumors.**

Representative confocal images of control (EJS) tumors, JNK-deficient EJS tumors (EJS + JNK^DN^), and JNK- and *Dronc-*deficient EJS tumors (EJS + JNK^DN^ + *Dronc*-i) after 1.5 days of EJS induction stained with anti-MMP1. Outline of wing disc (dashed white) obtained by tracing DAPI. Scale bar: 100μm.

**Supplementary Figure 3. JNK pathway activation is independent of ROS and JNK membrane receptors.**

**(A)** Representative confocal images of control (EJS) and EJS + UAS-*Dronc*-RNAi (EJS + *Dronc*-i) after 1 day of EJS induction labeled with the ROS indicator DHE. Inset shows the entire tumorous wing disc from which the main image (dashed outline) was magnified. Scale bar: 100μm.

**(B)**. Quantification of DHE levels in control (EJS) and EJS + UAS-*Dronc*-RNAi (EJS + *Dronc*-i) after 2 days of EJS induction, measured by counting the number of high-intensity DHE puncta per tumor. Statistical analysis performed by Student’s t-test. **** p<0.0001. Numbers of wing discs analyzed for EJS tumors: 7; for EJS + *Dronc*-i tumors: 7.

**(C)** Representative confocal images of control tumors (EJS), EJS + UAS-*Duox*-RNAi tumors (EJS + *Duox*-i), and EJS + UAS-*Catalase* tumors (EJS + Catalase) expressing GFP and hemocytes immunostained with anti-Hemese (H2). Notice the similar morphological features in all three conditions (GFP) and that hemocytes are still present even with the removal of ROS (H2). Scale bar: 100μm.

**(D).** Relative sizes of control EJS (EJS) and JNK receptor-deficient EJS tumors (EJS + *grnd-i* + *wgn-i*) after 3 days of EJS induction. Graphs show mean ±SD. Statistical analysis performed by an unpaired Student’s t-test. ns – not significant p>0.05. Numbers of wing discs analyzed for both EJS and EJS + *grnd-i* + *wgn-i* tumors: 49 each.

**Supplementary Figure 4. The overexpression of *reaper* in hemocytes of wild-type larvae without tumor causes a dramatic hemocyte ablation.**

Upper left: image of a *Drosophila* larva expressing GFP and Reaper specifically in hemocytes using the *hemolectin*-Gal4 (*hml*-G4) driver. No hemocytes can be detected. Lower right: in a sibling larva not expressing Reaper, hemocytes can be easily detected. Full genotype description in MM.

**GENOTYPE DESCRIPTION**

**Supplementary Figure 1:**

**(A and B)**

EJS: *ap-Gal4, UAS-GFP, UAS-Socs36E-RNAi/+; UAS-EGFR, tub-Gal80^ts^/+*

*EJS + Dronc-i: ap-Gal4, UAS-GFP, UAS-Socs36E-RNAi/Dronc-RNAi; UAS-EGFR, tub-Gal80^ts^/+*

**Supplementary Figure 2:**

WT: *w^1118^*

EJS: *ap-Gal4, UAS-GFP, UAS-Socs36E-RNAi/+; UAS-EGFR, tub-Gal80^ts^/+*

*EJS + Dronc-i: ap-Gal4, UAS-GFP, UAS-Socs36E-RNAi/Dronc-RNAi; UAS-EGFR, tub-Gal80^ts^/+*

*EJS + JNK^DN^: UAS-bsk^DN^; ap-Gal4, UAS-GFP, UAS-Socs36E-RNAi/+; UAS-EGFR, tub-Gal80^ts^/+*

**Supplementary Figure 3:**

**(A and B)**

EJS: *ap-Gal4, UAS-GFP, UAS-Socs36E-RNAi/+; UAS-EGFR, tub-Gal80^ts^/+*

*EJS + Dronc-i: ap-Gal4, UAS-GFP, UAS-Socs36E-RNAi/Dronc-RNAi; UAS-EGFR, tub-Gal80^ts^/+*

**(C)**

EJS: *ap-Gal4, UAS-GFP, UAS-Socs36E-RNAi/+; UAS-EGFR, tub-Gal80^ts^/+*

EJS + Duox-i: *ap-Gal4, UAS-GFP, UAS-Socs36E-RNAi/UAS-Duox-RNAi* (BL38907)*; UAS-EGFR, tub-Gal80^ts^/+*

EJS + Catalase: *ap-Gal4, UAS-GFP, UAS-Socs36E-RNAi/UAS-Catalase* (BL24621)*; UAS-EGFR, tub-Gal80^ts^/+*

**(D)**

EJS: *ap-Gal4, UAS-GFP, UAS-Socs36E-RNAi/+; UAS-EGFR, tub-Gal80^ts^/+*

EJS + grnd-I + wgn-i: *ap-Gal4, UAS-GFP, UAS-Socs36E-RNAi/UAS-Grindelwald-RNAi* (VDRC: 43454)*; UAS-EGFR, tub-Gal80^ts^/UAS-Wengen-RNAi* (BL50594)

**Supplementary Figure 4**

Hml>GFP, Reaper: *hml-QF2/QUAS-GFP* (BL52264)*; UAS-QS, QUAS-Reaper/+*

Hml>GFP: *hml-QF2/QUAS-GFP; TM6B/+*
