## Supplementary figures and images for "Widespread non-apoptotic activation of *Drosophila* Caspase-2/9 limits JNK signaling, macrophage proliferation, and growth of wound-like tumors"

### Supplementary Figure 1

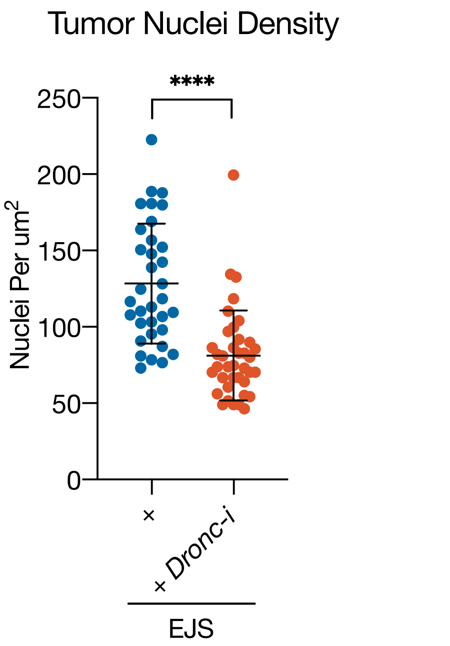


Xu et al., Supplementary Figure 1

### Supplementary Figure 2

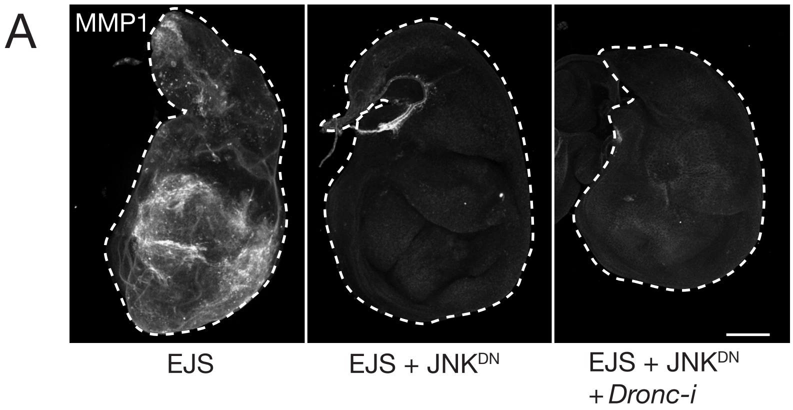


Xu et al., Supplementary Figure 2

### Supplementary Figure 3

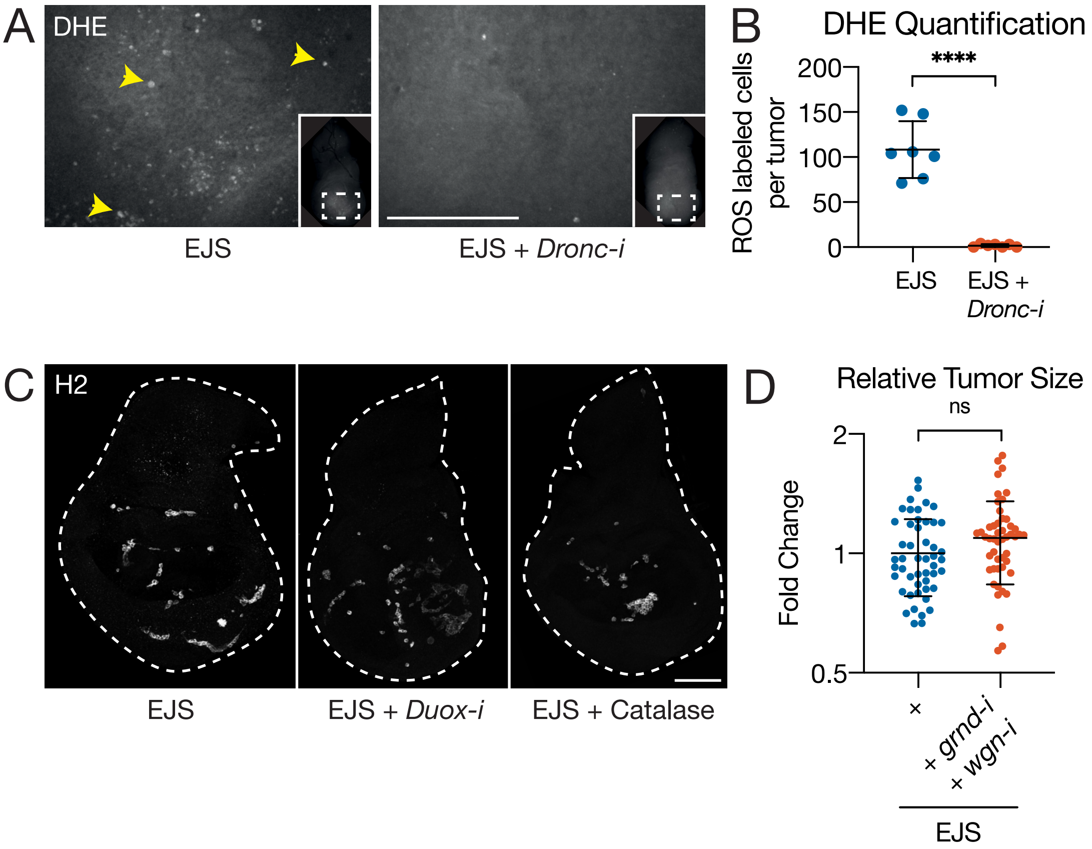


Xu et al., Supplementary Figure 3

### Supplementary Figure 4

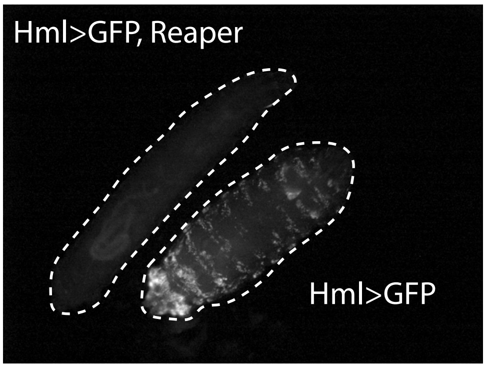


Xu et al., Supplementary Figure 4
